# Spatial-linked alignment tool (SLAT) for aligning heterogenous slices properly

**DOI:** 10.1101/2023.04.07.535976

**Authors:** Chen-Rui Xia, Zhi-Jie Cao, Xin-Ming Tu, Ge Gao

## Abstract

Spatially resolved omics technologies reveal the spatial organization of cells in various biological systems. Integrative and comparative analyses of spatial omics data depend on proper slice alignment, which should take both omics profiles and spatial context into account. Here we propose SLAT (Spatially-Linked Alignment Tool), a graph-based algorithm for efficient and effective alignment of spatial omics data. Adopting a graph adversarial matching strategy, SLAT is the first algorithm capable of aligning heterogenous spatial data across distinct technologies and modalities. Systematic benchmarks demonstrate SLAT’s superior precision, robustness, and speed vis à vis existing methods. Applications to multiple real-world datasets further show SLAT’s utility in enhancing cell-typing resolution, integrating multiple modalities for regulatory inference, and mapping fine-scale spatial-temporal changes during development. The full SLAT package is available at https://github.com/gao-lab/SLAT.

## Introduction

Recently emerging spatial omics technologies enable profiling the location, intercommunication, and functional cooperation of native cells through fluorescence *in situ* hybridization (seqFISH^1^, MERFISH^2^, seqFISH+^3^ and Xenium^4^) and spatial barcoding (10x Visium^5^, HDST^6^, Slide-seqV2^7^, Stereo-seq^8^ and spatial-ATAC-seq^9^) from multiple tissue “slices”, revealing tissue structure heterogeneity and shedding light on the underlying physiological and pathological mechanisms^8, 10^.

Properly aligning cells that share molecular features (e.g., cell type) and spatial context across multiple slices, especially these generated from distinct sources, is critical for their follow-up analysis. E.g., inter-technology alignment effectively bridges technologies complementary in spatial resolution and omics coverage^4^, while aligning various time points during spatially dynamic processes like embryogenesis helps identify key spatial temporal changes and their molecular underpinnings. However, current spatial alignment algorithms^11–13^ are mostly designed for homogeneous alignments (e.g., three-dimensional reconstruction from consecutive slices^14, 15^), and can hardly handle heterogeneous slices which often involve complex non-rigid deformations, uneven spatial resolutions as well as complex batch effects.

Here, we introduce SLAT (Spatially-Linked Alignment Tool), a unified framework for aligning both homogenous and heterogeneous single-cell spatial datasets. By modeling the intercellular relationship as a spatial graph, SLAT adopts graph neural networks and adversarial matching for robustly aligning spatial slices. In addition to its superior performance revealed by systematic benchmarks, as the first algorithm capable of aligning heterogenous spatial data, SLAT introduces a wide range of application scenarios including alignment across distinct technologies and experimental conditions. SLAT is publicly accessible at https://github.com/gao-lab/SLAT and will be continuously updated as spatial omics technologies evolve.

## Results

### Align heterogenous spatial omics data via graph adversarial matching

By modeling the spatial topology per slice as a spatial graph where each cell is connected to its nearest neighbors by edges, we reformulate the slice-alignment task as a graph-matching problem. In efforts to correct potential cross-dataset batch effect, SLAT employed an SVD-based matrix decomposition strategy to project omics profiles of the cells into a shared low-dimensional space (**Methods**), which in turn serves as node features of the spatial graphs.

Multilayer lightweight graph convolutional networks are incorporated to propagate and aggregate information between cells and their neighbors via stepwise concatenations, generating a holistic representation with information at multiple scales from individual cells to local niches as well as global positions. Then, SLAT solves a minimum-cost bipartite matching problem between the spatial graphs through a dedicated adversarial component^16^ to align cells from different slices (Fig. 1, **Methods**).

**Fig. 1.**
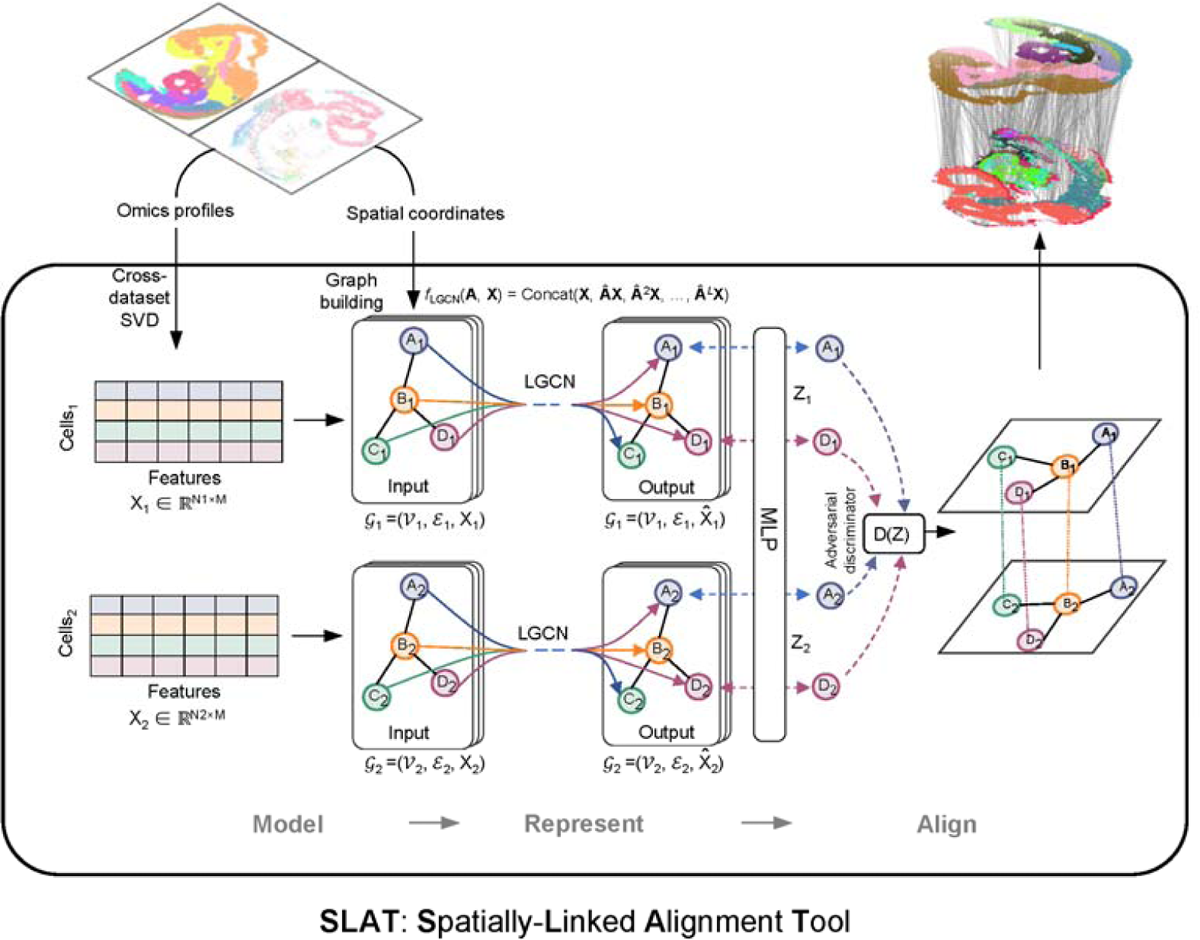
Architecture of SLAT framework. The SLAT algorithm can be divided into three main steps: model, represent and align. We model omics data from two single-cell spatial slices as low dimensional representations X_1_ ∈ R^N_1_x M^, X_2_ ∈ R^N_2_× M^ obtained via SVD-based cross-dataset decomposition, where *N*_1_, *N*_2_ are cell numbers of each slice and *M* is the SVD dimensionality. We use spatial coordinates of the cells to build *K*-neighbor spatial graphs g_1_ = (*V*_1_, ε_1_, X_1_), g_2_ = (*V*_2_, ε_2_, X_2_) in each slice, respectively. The neighbor size can be set dynamically to adjust for uneven spatial resolution across technologies. To encode local and global spatial information, SLAT uses *L*-layer lightweight graph convolution networks to propagate and aggregate information across the spatial graph into cell embeddings 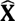_1_, 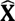_2_, which are then projected into alignment space z_1_, z_2_ with a multi-layer perceptron trained adversarially against a discriminator *f*_D_ (z) to align the two graphs by minimizing their Wasserstein distance (see **Methods**). Node labels with the same letters (e.g., *A*_1_, *A*_2_) represent corresponding cells in different slices.

Notably, apart from congruent regions that can be well aligned across slices, heterogeneous alignment often involves distinct regions that reflect biologically relevant spatial alterations. To avoid artificially aligning such regions (i.e., over-alignment), SLAT adopts an adaptive clipping strategy during adversarial matching to retain only the closest pairs between slices in terms of their cosine similarity in the embedding space, taken as reliable anchors for guiding the whole alignment procedure (Extended Data Fig. 1a). Furthermore, SLAT calculates a similarity score for each aligned pair, making it possible to pinpoint spatially discrepant regions (see below and **Methods** for more details).

### Systematic benchmarks suggest SLAT is accurate robust and fast

To evaluate SLAT’s performance over existing algorithms which are designed for homogeneous alignment^12, 14^, we first ran systematic benchmarks based on consecutive, homogeneous slices from the same tissue generated by three representative technologies with a wide range of throughput, resolution, and technological routes: 10x Visium^5^, MERFISH^2^, and Stereo-seq^8^ (Fig. 2a, Supplementary Data Table 1, Supplementary Fig. 1). All slices were randomly rotated before feeding to the alignment methods, which in turn attempt to recover the correct cell matching in expert curated cell types and spatial regions (as visualized by vertical lines connecting the two slices in Fig. 2b and as heatmaps in Fig. 2c).

**Fig. 2.**
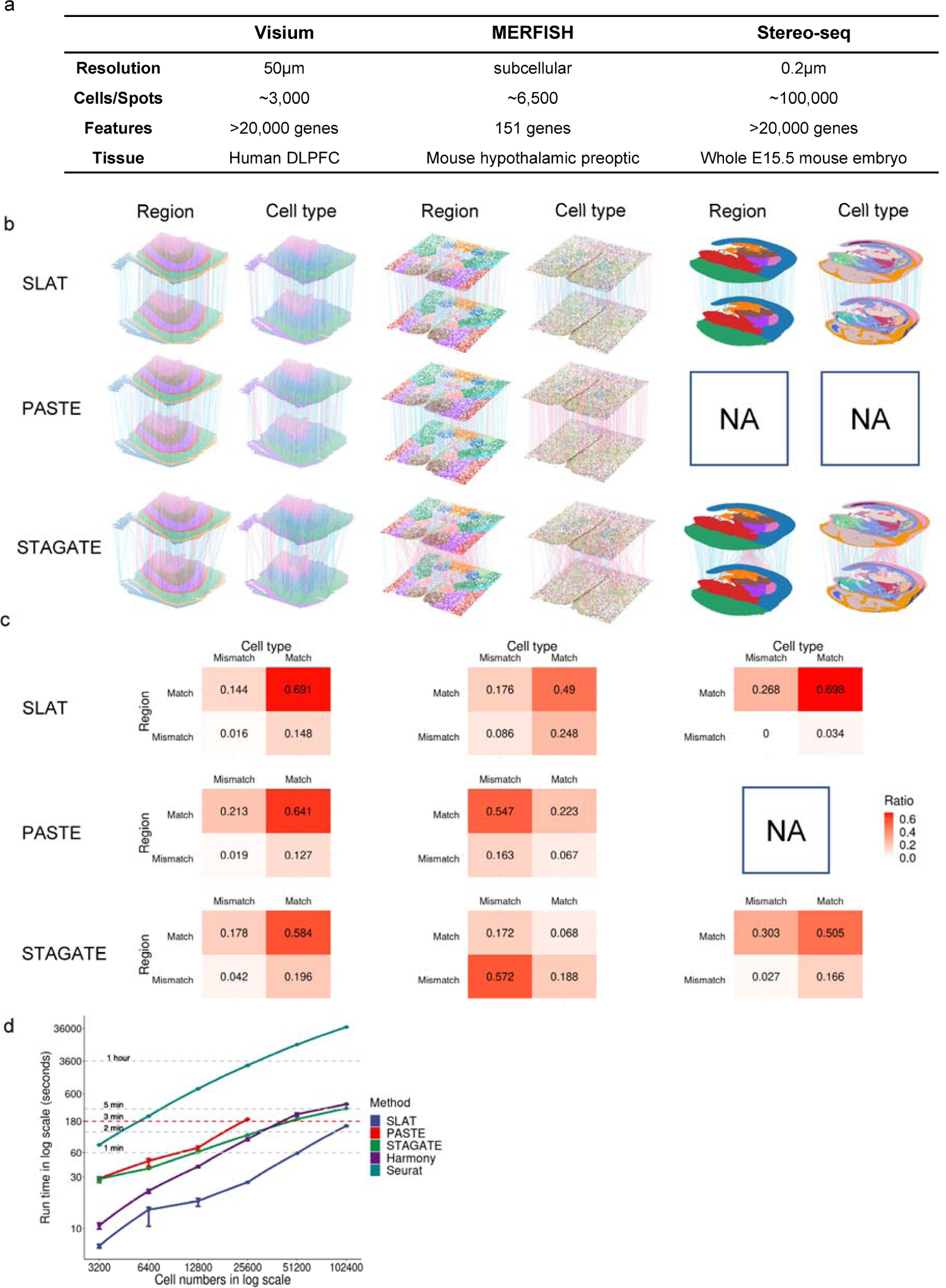
Evaluation on homogeneous spatial alignment. **a**, Summary of the three benchmark datasets. For each dataset, two slices were used for alignment, see Supplementary Fig. 1 for more details of every slice. **b**, Visualization of alignment results of different methods on the benchmark datasets in **a**. Slices are colored by spatial region (left) and cell types (right), respectively. Vertical lines connect aligned cell pairs (subsampled to 300 alignment pairs for clear visualization). Blue lines indicate accurate alignments (with matching cell types and spatial regions) while red lines indicate inaccurate alignments. PASTE failed to run on the Stereo-seq dataset due to GPU memory overflow (capping at 80 GB). **c**, Heatmaps quantifying the region matching accuracy and cell type matching accuracy respectively in **b**, in the form of confusion matrices. The number in each cell is the average proportion across eight repeats with different random seeds. **d**, Running time of each method on subsampled datasets of varying sizes. *n* = 8 repeats with different subsampling random seeds. Error bars indicate mean ± s.d.

Overall, SLAT outperformed PASTE^14^ and STAGATE^12^ in all three datasets (Fig. 2b and c). Of note, PASTE exhibited particularly low cell type accuracy in the MERFISH dataset where different cell types are spatially interlaced (Fig. 2c), probably due to its excessive reliance on the spatial distance between cells over molecular features. Meanwhile, we found that STAGATE produced suboptimal alignments albeit employing graph neural networks for modeling spatial omics data too, while its performance improved for split slices which are free of batch effects (**Methods**, Supplementary Fig. 2).

By combining spatial context with transcriptome profiles, SLAT is better equipped to distinguish transcriptionally similar but spatially distinct cell groups. Consistently, we found that SLAT also achieved significantly higher alignment accuracy than the conventional spatially unaware algorithms Seurat^17^ and Harmony^18^ (Supplementary Fig. 2-5). In particular, although these algorithms matched cell types reasonably well, their alignment largely disarranged spatial regions (Supplementary Fig. 2-5).

SLAT’s lightweight graph convolutional component effectively improves model robustness. Evaluation with subsampled data of various sizes showed that SLAT consistently provides the best results, even with as few as 200 cells (Supplementary Fig. 5, **Methods**). Further inspection confirmed that the performance of SLAT remains highly robust to a wide range of hyperparameter settings and against random corruption of the spatial graph (Extended Data Fig. 2, **Methods**).

As technologies continue to evolve, the throughput of spatial single-cell experiments is constantly increasing^19^. Implemented as a neural network and optimized for parallelism, SLAT is highly scalable: benchmarks showed that SLAT is consistently the fastest method across all data sizes and aligns slices each with over 100,000 cells in just 3 minutes (Fig. 2d). We also noticed that PASTE fails to run as cell number exceeds 25,600, partly due to its memory intensive implementation for optimal transport.

### Matching heterogeneous datasets across distinct technologies and modalities

Spatial alignment across multiple technologies and modalities amalgamates complementary information towards a comprehensive *in situ* view of cellular states^20^. Benefiting from its novel design, SLAT provides reliable alignment across heterogeneous datasets, which is largely beyond the capability of current alignment algorithms.

We first used SLAT to align spatial transcriptomic datasets generated by two distinct technologies with varying scale and detectability^8, 21^. Specifically, the seqFISH dataset contains over 10,000 cells with only 350 genes detected in total^3, 21^, whereas the Stereo-seq dataset contains about 5,000 cells covering over 20,000 genes^8^. Meanwhile, the cell types in the two datasets were annotated at different resolutions: the seqFISH dataset has finer-grained annotation (21 cell types, Extended Data Fig. 3a) than the Stereo-seq dataset (11 cell types, Extended Data Fig. 3b). The high-quality alignment produced by SLAT (Fig. 3a) enables an accurate label transfer for cell typing improvement (Extended Data Fig. 3c). For example, cells labeled as “Neural crest” in the Stereo-seq dataset were aligned to seqFISH regions with four major cell types (Fig. 3b, Extended Data Fig. 3d), refining cell typing which was further validated by known marker genes (Fig. 3c and Extended Data Fig. 3e).

**Fig. 3.**
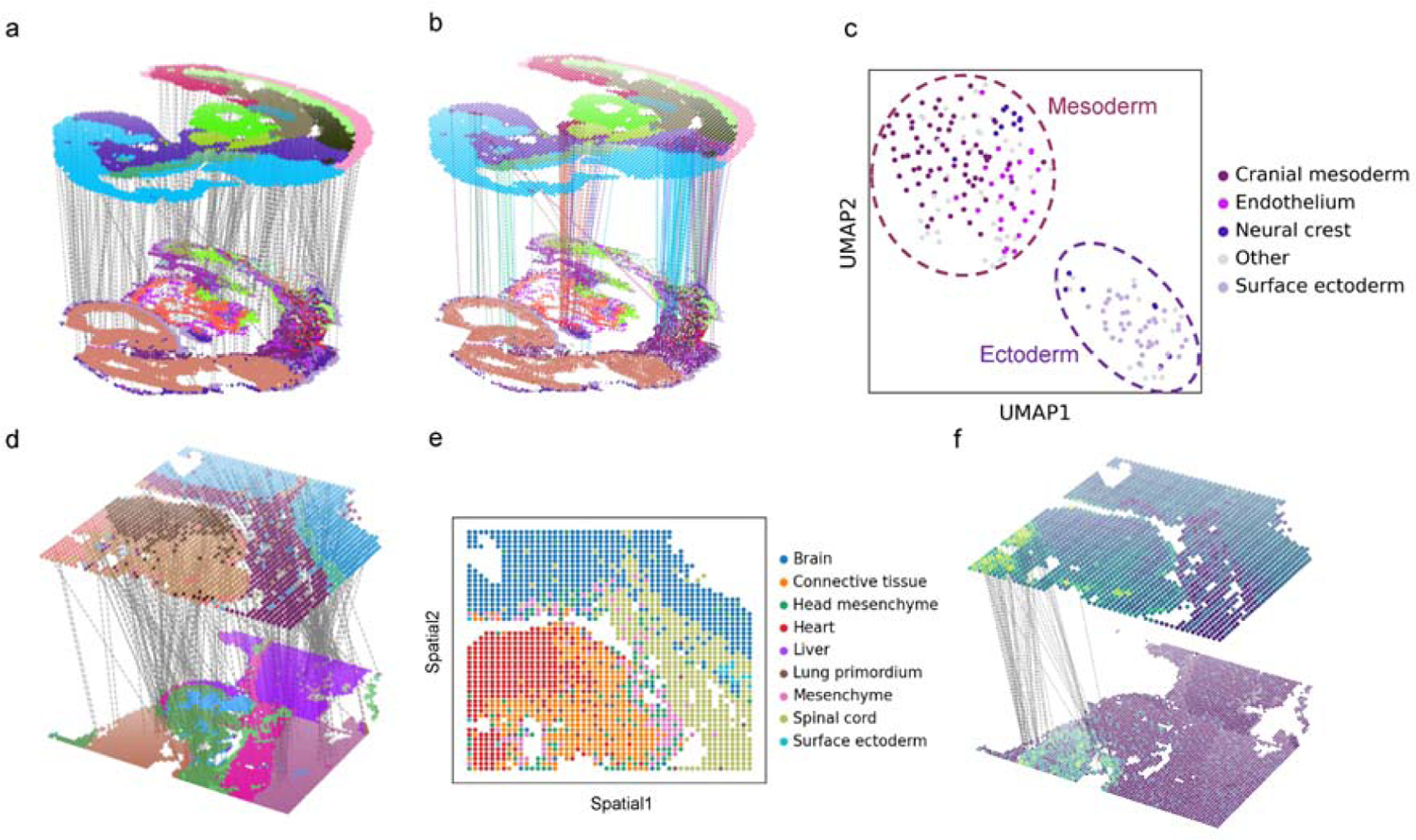
Spatial alignment across distinct technologies and modalities. **a**, Visualization of the alignment of E8.75 seqFISH mouse embryo and E9.5 Stereo-seq mouse embryo datasets, colored by cell types (subsampled to 300 alignment pairs for clear visualization). **b**, Highlighting the alignment of cells labeled as “Neural crest” in Stereo-seq (top) to seqFISH (bottom), alignment lines are colored by cell type of aligned cells in seqFISH. **c**, UMAP visualization of cells labeled as “Neural crest” in Stereo-seq, colored by cell type of aligned cells in seqFISH, cell types with fractions less than 5% are collectively labeled as “Other”. Dashed circles indicate manual annotation by marker genes (corresponding to Extended Data Fig. 3c and 3d). **d**, Visualization of the alignment of E11.5 spatial-ATAC-seq mouse embryo and E11.5 Stereo-seq mouse embryo datasets, with spatial-ATAC-seq dataset colored by clusters (see Extended Data Fig. 6e) and Stereo-seq colored by cell types, respectively (subsampled to 300 alignment pairs for clear visualization). **e**, Cell type labels of spatial-ATAC-seq transferred from Stereo-seq via SLAT alignment. **f**, Showing chromatin accessibility score and gene expression pattern of heart marker *Tnnt2* on SLAT alignment.

Notably, slices in the two datasets also differ in spatial structure with substantial non-rigid deformations (Extended Data Fig. 3a and 3b). Such challenging scenario effectively failed methods other than SLAT (Extended Data Fig. 4). Similar cross-scale alignment can also be achieved on Visium and Xenium slices^4^, where SLAT accurately pinpoints a rare group of triple positive breast tumor cells, while all other methods failed (Extended Data Fig. 5).

**Fig. 4.**
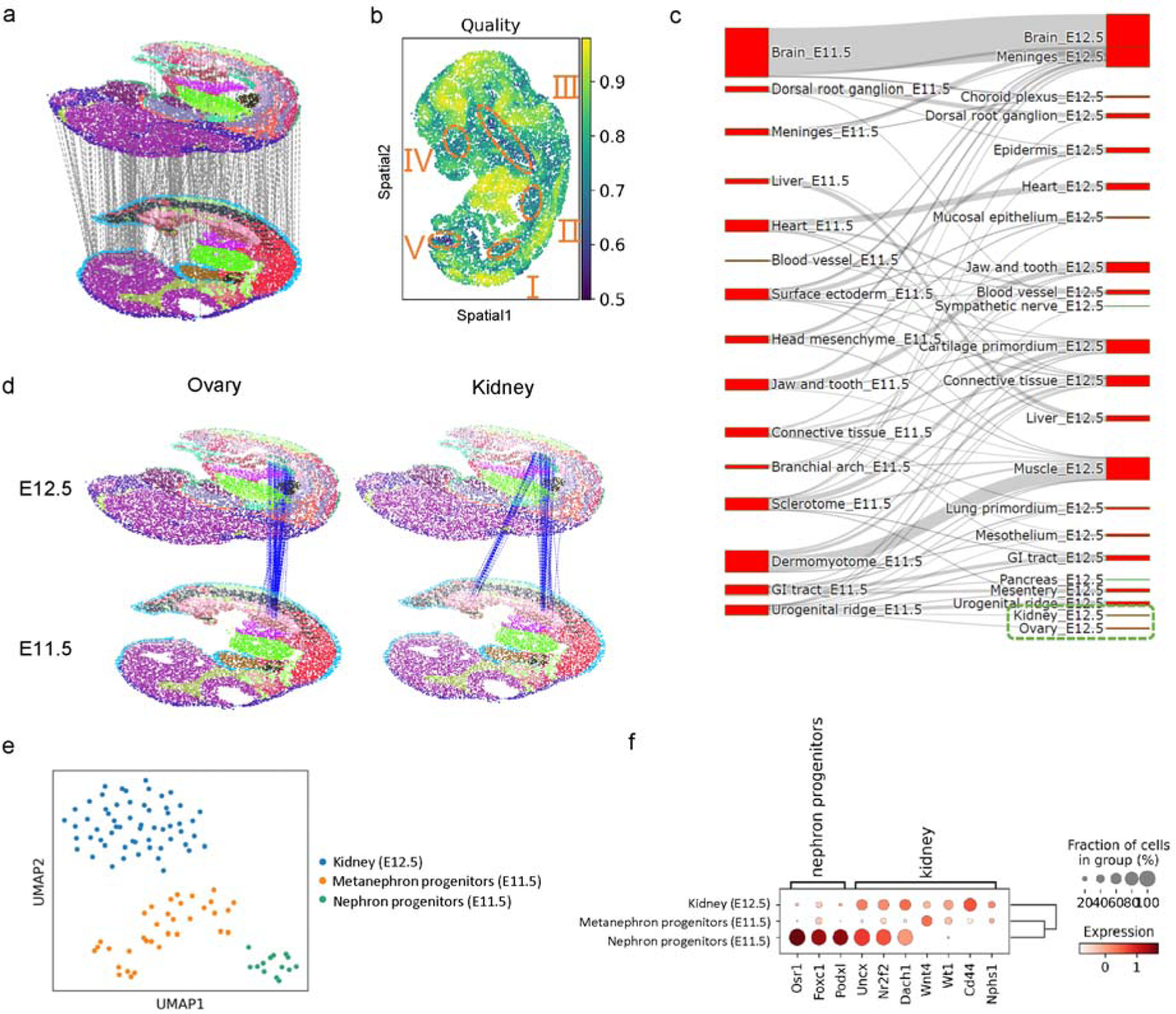
Developmental alignment of mouse embryo. **a**, Visualization of the alignment of E11.5 and E12.5 mouse embryo Stereo-seq datasets, colored by cell types (subsampled to 300 alignment pairs for clear visualization). **b**, Similarity score of the alignment. Higher scores indicate higher alignment confidence. Dashed circles highlight five regions with low similarity scores. **c**, Sanky plot showing cell type correspondence of SLAT alignment between the two datasets. The green box highlights cells labeled as “Kidney” and “Ovary” in E12.5. **d**, Alignment visualization highlighting cells labeled as “Ovary” (left) and “Kidney” (right) in E12.5 and their aligned cells in E11.5, respectively. **e**, UMAP visualization of cell type annotations for cells labeled as “Kidney” in E12.5 and their aligned cells in E11.5. **f**, Dot plot showing marker gene expression of cell types in **e**.

More challenging is the cross-modality spatial alignment, which is partly due to the disjoint feature spaces that invalidate the use of our SVD-based cross dataset matrix decomposition strategy as well as other canonical batch correction methods. Benefitting from the modular design of SLAT (Fig. 1), we employed the graph-linked multi-modality embedding strategy we proposed before^22^ to project cells of different modalities into a shared embedding space before feeding them into the LGCNs (**Methods**). With this extension, SLAT successfully produced a spatial alignment across RNA (Stereo-seq) and ATAC (spatial-ATAC-seq) slices. While the different modalities featured drastically different spatial resolutions (0.2 μm for Stereo-seq but 20 μm for spatial-ATAC-seq), SLAT managed to align them well (Fig. 3d, Extended Data Fig. 6a). Cell-type labels transferred from Stereo-seq to spatial-ATAC-seq based on the alignment were consistent with anatomical features and with the accessibility of tissue-specific genes (Fig. 3e and 3f, Extended Data Fig. 6b). Joint regulatory analysis using the aligned cell pairs and transferred labels effectively identified various key regulators in the heart, such as *Jund* and *Ctnnb1*^23, 24^ (**Methods**, Extended Data Fig. 6c). We also experimented with other spatial alignment methods by feeding the same multi-modality embeddings as input, but found suboptimal results (Supplementary Fig. 7).

### Mapping fine-scale spatial-temporal transitions by developmental alignment

Embryonic development is a highly dynamic process with extensive spatial-temporal transitions involving the generation, maturation, and functional alteration of organs and tissues at specific timepoints. To probe spatial-temporal dynamics during early development, we used SLAT to align two spatial atlases of mouse embryonic development at E11.5 and E12.5^8^ (Fig. 4a).

Most cells in the brain, heart, and liver were well aligned with high alignment credibility scores, consistent with their spatial-temporal conservation at the two timepoints (Fig. 4b, Extended Data Fig. 7a and 7b). In addition, several regions were enriched with less-aligned cells of lower alignment credibility scores, which may be attributed to both biological development (e.g., the newly emerged organs kidney and ovary at region I^8, 25–27^, the rapid enlargement of lung primordium and its displacement from the upper part of the heart to the lower part at region II^28^ as shown in Extended Data Fig. 7c, and the disappearance of branchial arch at corresponding position of region III in E11.5^8^) and technical variation (e.g., tissue loss at corresponding position of region IV in E11.5).

We next followed up on kidney and ovary, two newly emerging organs at around E11.5–12.5^8, 25–27^. SLAT accurately identified them both as developing from the “Urogenital ridge” at E11.5 (highlighted by the green box in Fig. 4c). Consistent with previous reports, SLAT alignment showed precisely that the ovary develops directly from a single area^29^ (left panel of Fig. 4d) while the kidney develops from two separate areas corresponding to the mesonephros and metanephros structures in early kidney development^25, 30, 31^ (right panel of Fig. 4d, Extended Data Fig. 7d). In addition, based on the clustering analysis of aligned cells, we further identified a group of rare nephron progenitor cells located in the mesonephros, likely corresponding to an ephemeral mesonephric tubule during mesonephros development^32^ (Fig. 4e and 4f). Interestingly, the results show that the mesonephros is spatially adjacent to the ovary, which is consistent with the well-documented developmental cascade whereby a part of the mesonephros develops into the fallopian tube after degeneration (Fig. 4d). Our findings revealed the unique value of heterogeneous spatial alignment for interrogating the spatial-temporal process of organogenesis.

We also attempted the same alignment task with other methods (Extended Data Fig. 8). The spatially unaware algorithms (Seurat, Harmony) simply missed mesonephros, while the spatially aware algorithms (PASTE, STAGATE) align them to incorrect cells, further demonstrating the unique capability of SLAT on heterogeneous spatial alignment.

## Discussion

One of the essential challenges in spatial omics alignment is to appropriately model spatial context. Early methods such as Splotch^13^ and Eggplant^33^ model slices as rigid bodies and require manual annotation of landmark spots to guide the alignment. PASTE^14^ eliminates the need for landmark annotation by considering gene expression of all spatial spots, but its reliance on exact spatial distance impedes application to heterogeneous alignments involving complex non-rigid structural alterations. By combining spatial graph convolution and adversarial matching, SLAT achieves reliable spatial alignment for both homogeneous and heterogeneous slices in an unsupervised, data-oriented manner.

3D reconstruction from consecutive slices is a common application of spatial alignment. SLAT supposes reconstruction from multiple slices via progressive pairwise alignment (e.g., Extended Data Fig. 9) for a consecutive stack of four slices from the same E15.5 mouse embryo). While similar 3D stacking can also be achieved with PASTE^14^, SLAT is better equipped to account for non-rigid structural shifting and alteration among slices, enabling adaptive correction for potential deformation artifacts (see Supplementary Fig. 9 for quantitative assessment).

SLAT’s unique ability to conduct heterogeneous spatial alignments promises a wide range of biological applications. In particular, such spatial alignment enables identifying spatially-resolved changes such as key alterations of spatial patterns during development. Meanwhile, proper cross-modal alignment further sheds lights on key regulators and corresponding regulatory circuits. However, it should be noted that the alignments produced by SLAT do not necessarily presuppose one-to-one matchings: the best match returned by SLAT should be regarded as the cell with the highest similarity in terms of cell type and spatial context, and should not be taken as definite one-to-one matches as in single-cell multi-omics techniques where different modalities were generated from the same cell simultaneously.

SLAT is fast. In fact, generating the input cell embeddings is the most time-consuming step, while training and inferring with the SLAT core model takes only about 10 seconds for 10^6^ cells. Once the input embeddings are ready, aligning millions of cells can be completed in near real-time, enabling an efficient search of a massive database within an affordable timeframe. SLAT’s blazing speed sets the stage for some exciting applications such as inferring spatially dependent causal mechanisms by systematically comparing multiple Perturb-map slices^34^, or constructing whole organ 3D atlases involving thousands of slices^35^.

Last but not least, designed as a flexible framework, SLAT can be readily adapted and extended. For instance, additional information such as expert curation may be incorporated into the coordinate matching module to help distinguish symmetric structures and polish the final alignment (**Methods**). Meanwhile, the spatial graph modeling technique employed in SLAT may also be adapted to address other problems, such as comparative alignment across species.

Overall, SLAT provides a unified framework for various spatial integration scenarios. To promote its application by the research community, the SLAT package, along with detailed tutorials and demo cases, is available online at https://github.com/gao-lab/SLAT.

## Acknowledgments

We thank Mr. H. Yang for his helpful guidance on mouse embryonic development and anatomy, as well as Drs. Z. Zhang, L. Tao, F. Tang, X.S. Xie, C. Li, and J. Lu at Peking University for their helpful discussions and comments during the study. This work was supported by funds from the National Key Research and Development Program of China (2022ZD0115004), as well as the State Key Laboratory of Protein and Plant Gene Research, the Beijing Advanced Innovation Center for Genomics (ICG) at Peking University, the Changping Laboratory, and the Shaw Foundation Hong Kong Limited. The research of G.G. was supported in part by the National Program for Support of Top-Notch Young Professionals.

## Author contributions

G.G. and Z.J.C conceived the study and supervised the research. C.R.X and Z.J.C. designed and implemented the computational framework and conducted benchmarks and case studies with guidance from G.G. X.M.T proposed and implemented the matrix decomposition-based batch correction strategy. C.R.X, Z.J.C., X.M.T and G.G. wrote the manuscript.

## Extended Data Figures

**Extended Data Fig. 1.**
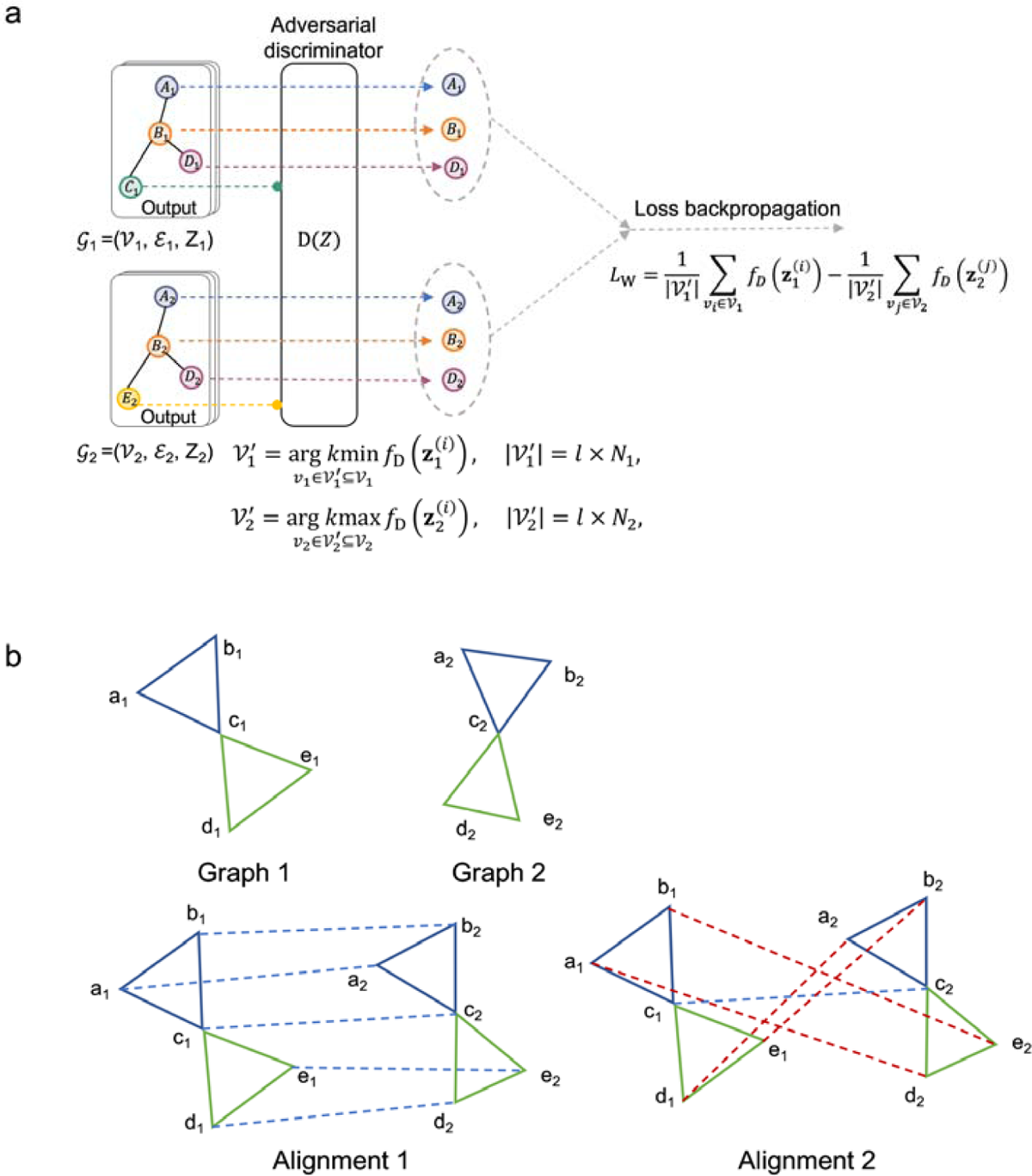
Illustration of dynamic clipping and alternative metrics. **a**, Illustration of the dynamic clipping criteria used to select cells for adversarial training. Only a subset of cells from the two datasets with minimal Wasserstein distance are selected (see **Methods**). **b**, Illustration where the “edge score” metric is ineffective. “Graph 1” and “Graph 2” are two graphs with five nodes, where nodes with the same letters (e.g., a_1_, a_2_) represent ground truth node pairs. “Alignment 1” and “Alignment 2” are two graph-matchings: “Alignment 1” correctly matches all nodes between two graphs, while only one node is correctly matched in “Alignment 2”. Nevertheless, both alignments give the same “edge score”.

**Extended Data Fig. 2.**
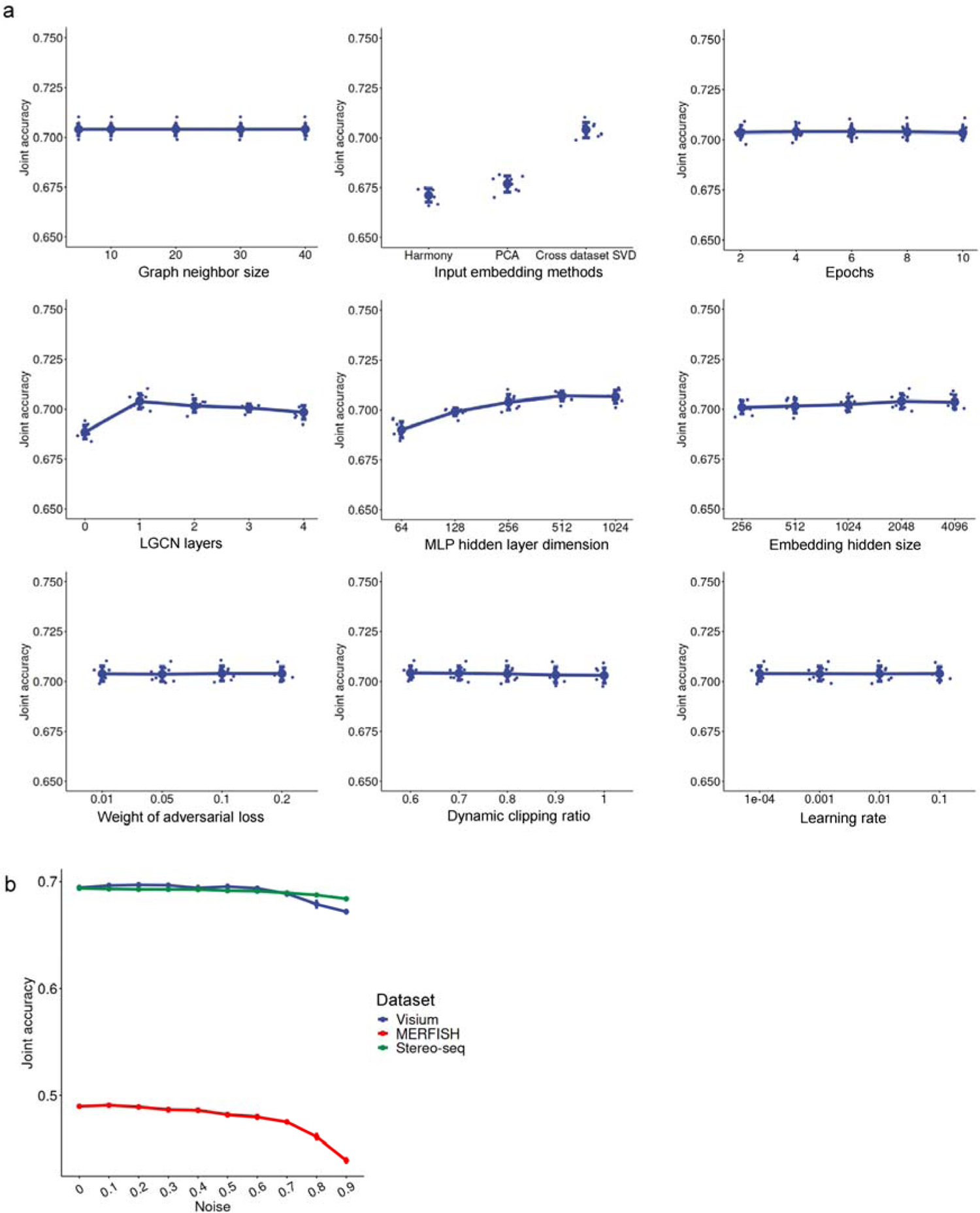
Effect of hyperparameter settings on alignment performance. **a**, Joint accuracy, which is defined as the proportion of aligned cells with both cell type and spatial region matched correctly (**Methods**) for SLAT with different hyperparameter settings including: graph neighbor size (K), input embedding methods, number of training epochs, number of LGCN layers (L), MLP hidden layer dimension, embedding hidden size (P), weight of the adversarial loss (α), dynamic clipping ratio (l), and learning rate. *n* = 8 repeats with different model random seeds. We scaled the *y*-axis to [0.65, 0.75] to show subtle changes. See **Methods** for detailed explanations of the hyperparameters. **b**, Changes in joint accuracy at different spatial graph corruption rates. *n* = 8 repeats with different model random seeds.

**Extended Data Fig. 3.**
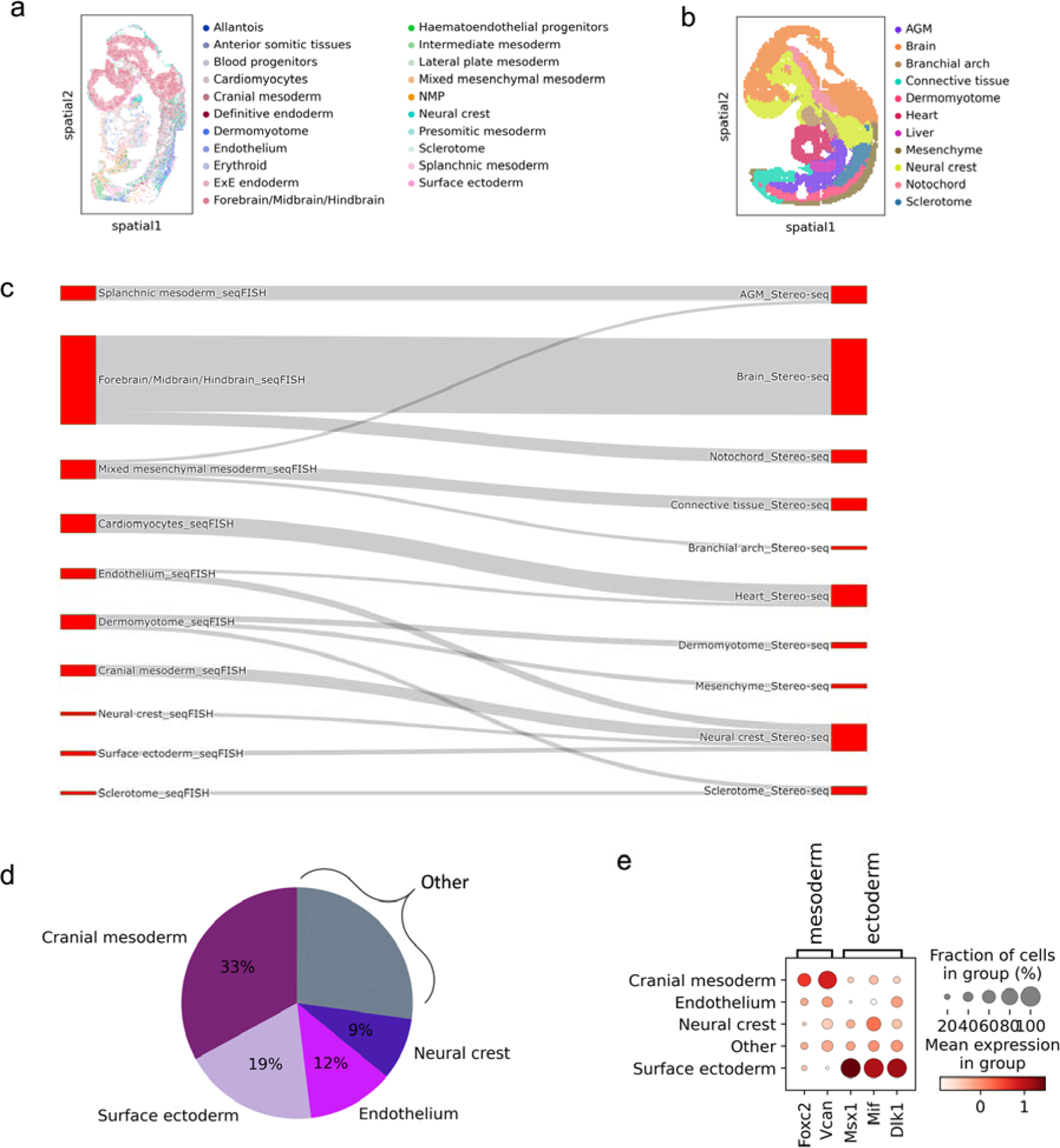
Refining cell type annotation in Stereo-seq via label transfer. **a-b**, Original annotation of **a**, the seqFISH dataset and **b**, the Stereo-seq dataset. **c**, Sanky plot showing cell type correspondence of SLAT alignment between seqFISH and Stereo-seq dataset. **d**, Proportions seqFISH transferred cell types in Stereo-seq cells originally labeled as “Neural crest”. All other cell types with proportions of less than 5% are collectively shown as “Other”. **e**, Expression of ectoderm and mesoderm marker genes as partitioned in Fig. 5c.

**Extended Data Fig. 4.**
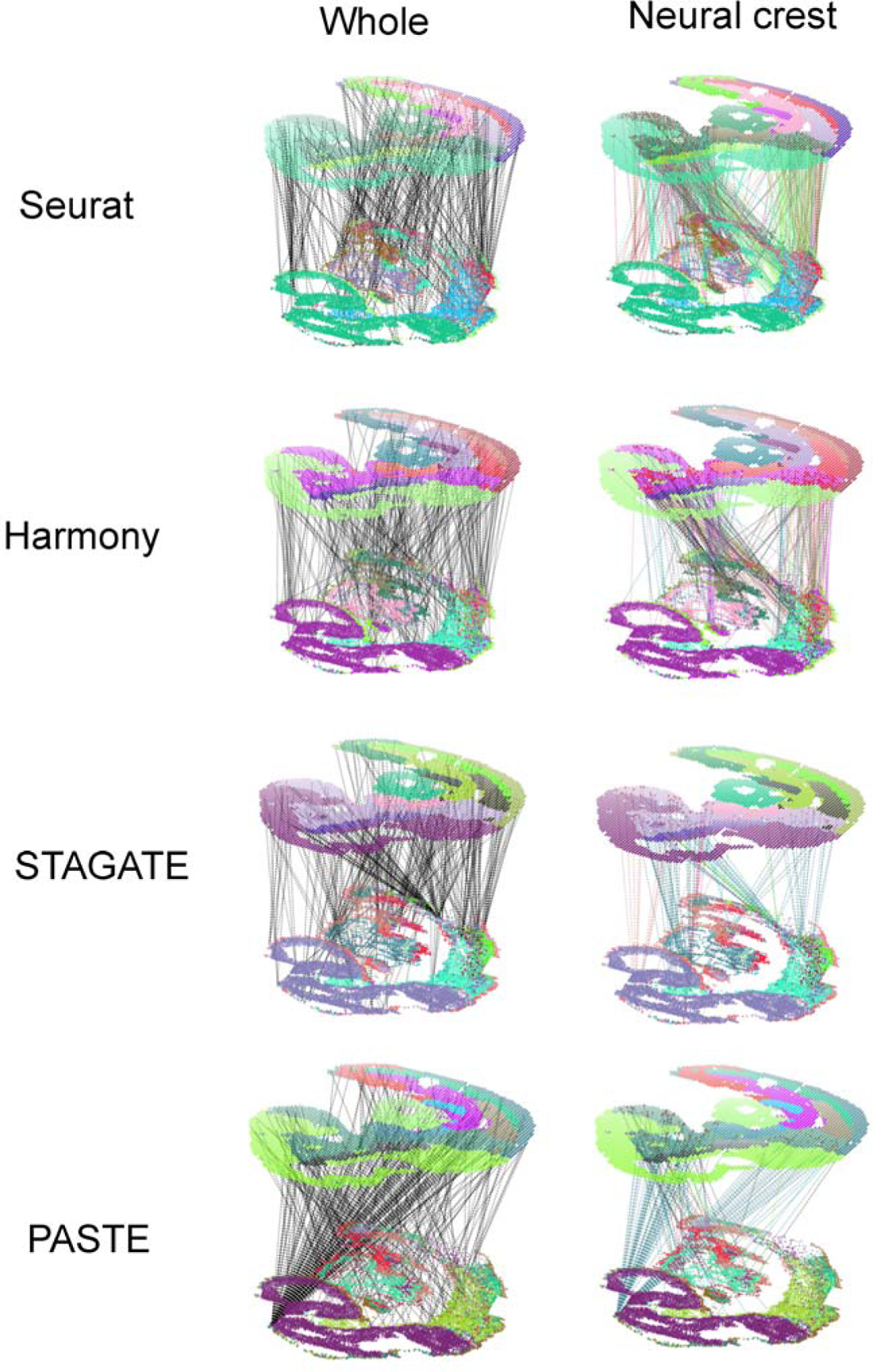
Alignment results of current methods on the seqFISH and Stereo-seq slices. Visualization of the alignment results of seqFISH and Stereo-seq mouse embryo slices via current methods. The left panels show the complete alignment results (subsampled to 300 alignment pairs for clear visualization), while the right panels highlight cells labeled as “Neural crest” in Stereo-seq and their aligned cells in seqFISH.

**Extended Data Fig. 5.**
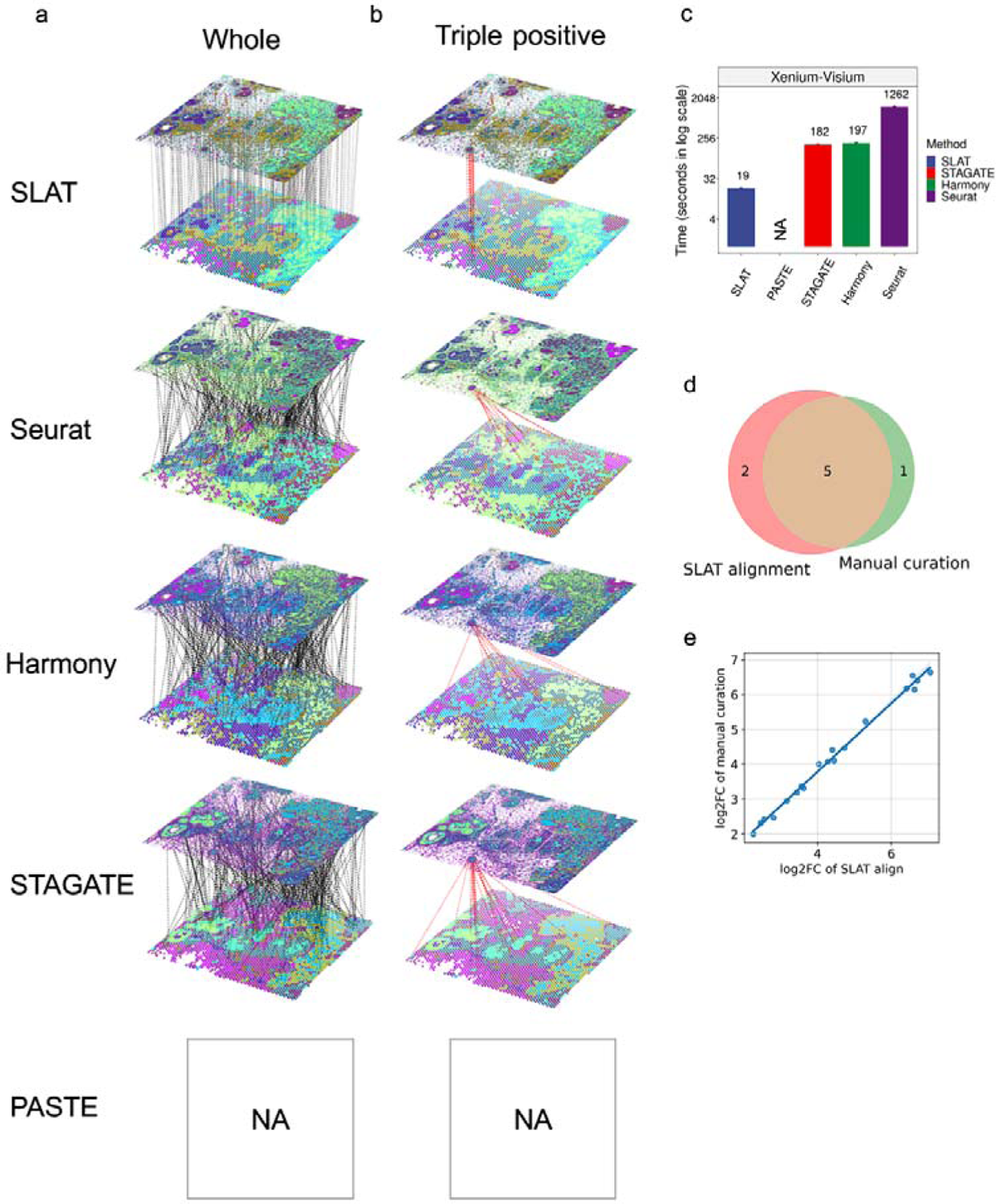
Alignment results of current methods on the Xenium and Visium slices. Xenium and Visium are spatial technologies designed by 10x with complementary capabilities^4^. Xenium achieves subcellular resolution but only detects a limited set of genes, while Visium profiles the complete transcriptome with coarse spatial resolution. Specifically, the Xenium slice covers more than 100,000 cells with 313 genes detected in total, and the Visium slice contains 3,841 spots with over 20,000 genes^4^. **a**, **b**, Visualization of the alignment results of two consecutive Xenium and Visium human breast cancer slices via different methods (subsampled to 300 alignment pairs for clear visualization), **b** Highlighting cells labeled as “Triple positive breast tumor cells” (ERBB2+/ESR1+/PGR+, Supplementary Fig. 6a and 6b) in Xenium and their aligned cells in Visium. PASTE failed due to GPU memory overflow (capping at 80 G). Of note, SLAT is the only algorithm that successfully aligned Xenium-resolved rare triple positive cells to Visium slice, even though they span less than 10 spots (Supplementary Fig. 6c and 6d). **c**, Running time of different methods. **d**, Overlapping of spots between the SLAT aligned and manually curated triple positive spots. **e**, Log_2_ fold change of shared top 25 marker genes in **e** for SLAT aligned compared to manually curated triple positive spots (Supplementary Fig. 6e).

**Extended Data Fig. 6.**
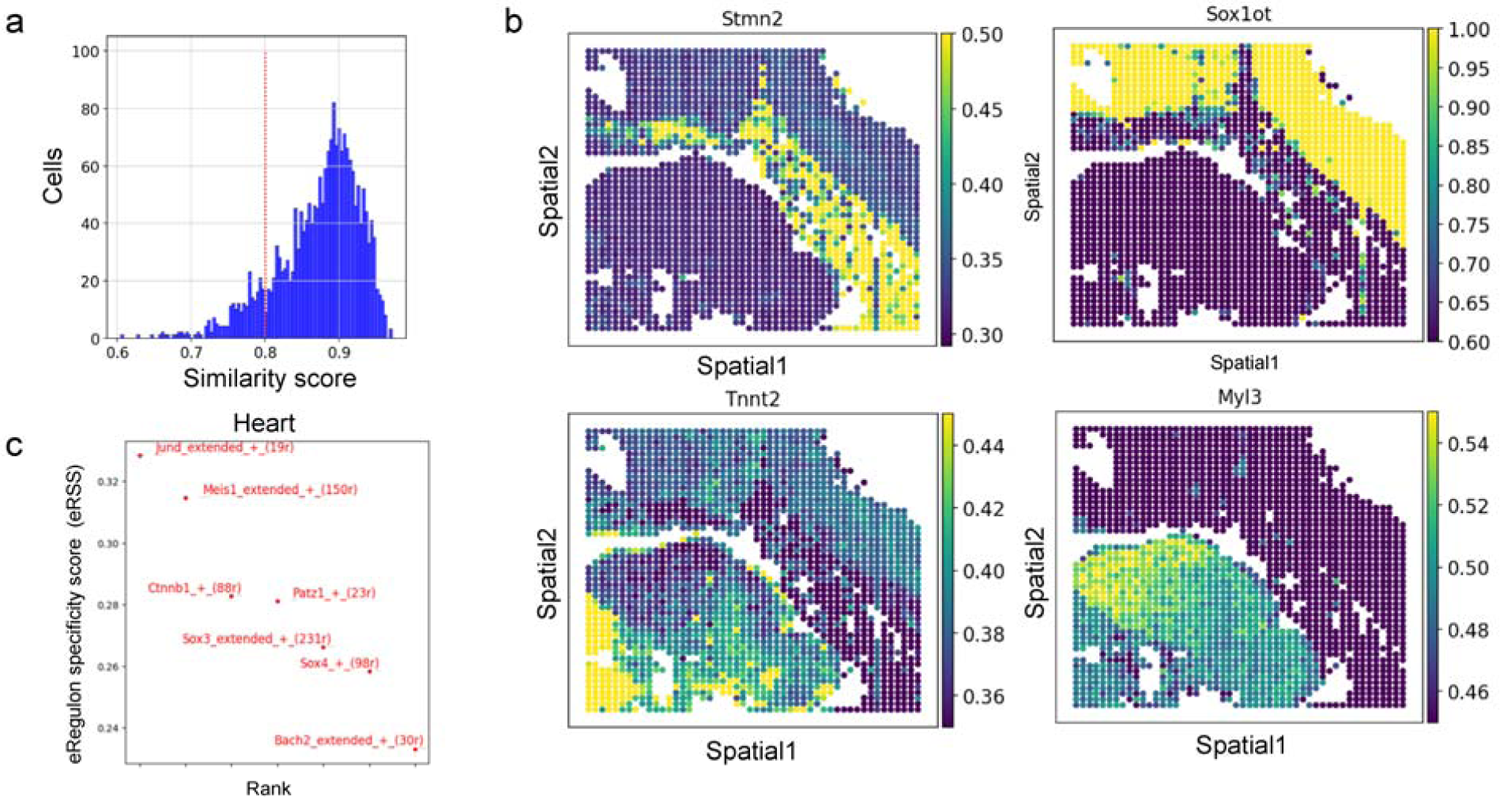
Alignment-based cross-modality annotation and regulatory reference. **a**, Similarity score distribution in spatial-ATAC-seq and Stereo-seq alignment. **b**, Visualization of chromatin accessibility score of cell type marker genes in the spatial-ATAC-seq slice. **c**, Key heart regulators identified by SCENIC+ based on the SLAT alignment.

**Extended Data Fig. 7.**
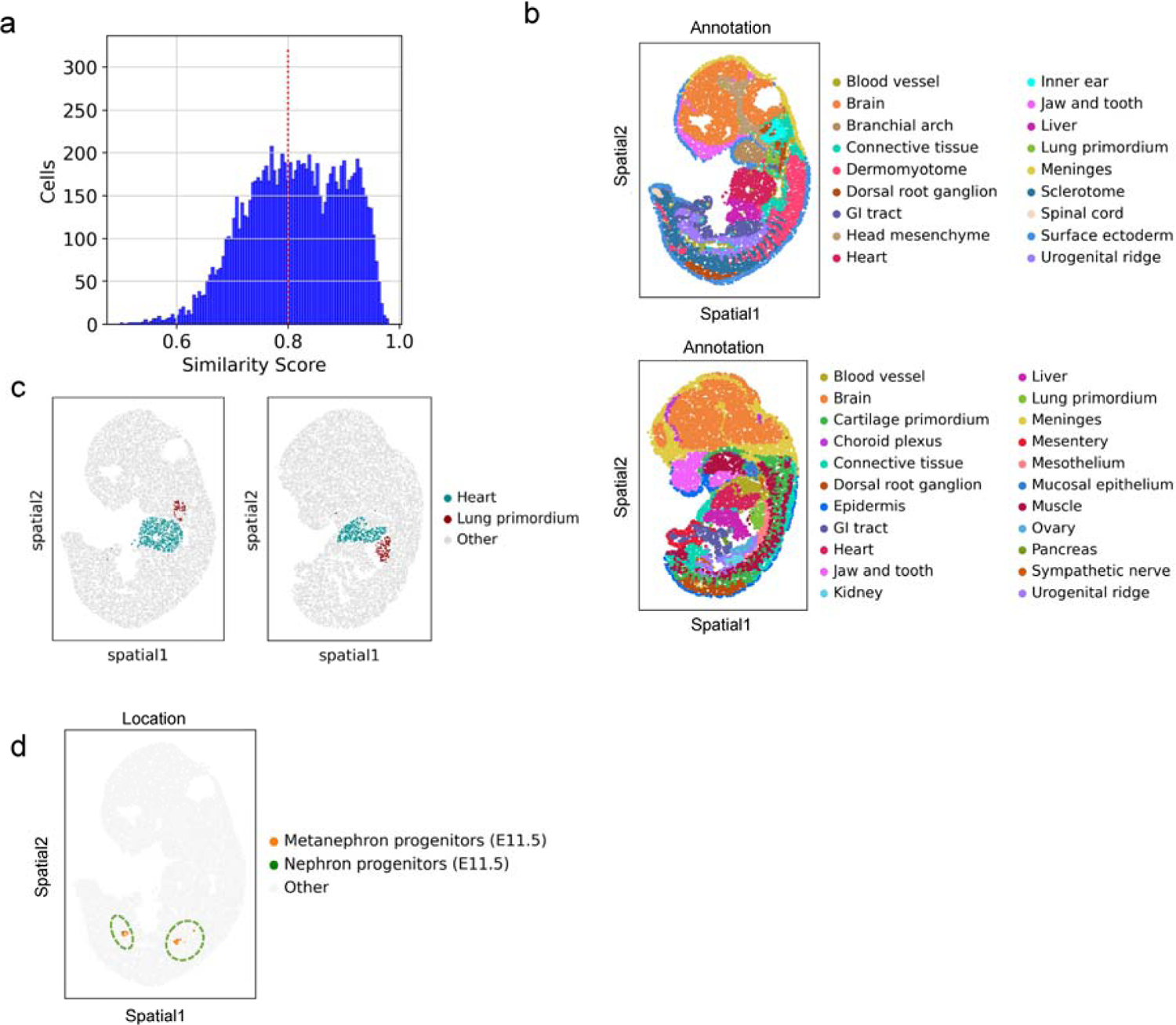
Spatial-temporal dynamics in mouse embryonic development. **a**, Similarity score distribution in E11.5 and E12.5 mouse embryo Stereo-seq slice alignment. **b**, Original annotation of E11.5 (top) and E12.5 (bottom) mouse embryo datasets. **c**, Spatial location of “Heart” and “Lung primordium” cells in the E11.5 (left) and E12.5 (right) slice, respectively. **d**, Spatial location of “Kidney” aligned cells in E11.5 in the right panel of **d**, colored by cell type in Fig. 4**e**.

**Extended Data Fig. 8.**
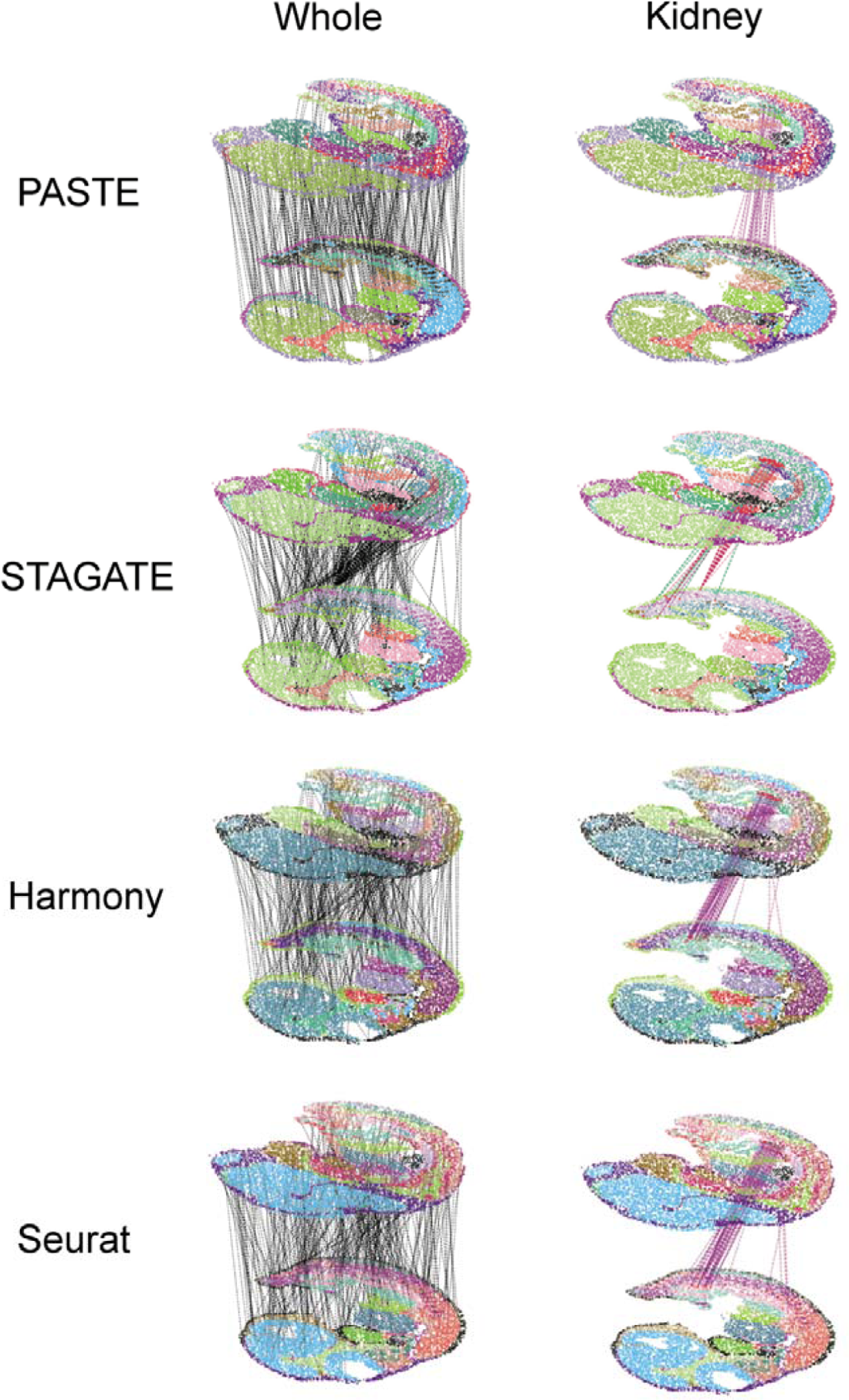
Alignment results of current methods in mouse embryonic development. Alignment of E11.5 and E12.5 mouse embryo slices using current methods. The left panels show the complete alignments (subsampled to 300 alignment pairs for clear visualization), while the right panels highlight cells labeled as “Kidney” in E12.5 and their aligned cells in E11.5.

**Extended Data Fig. 9.**
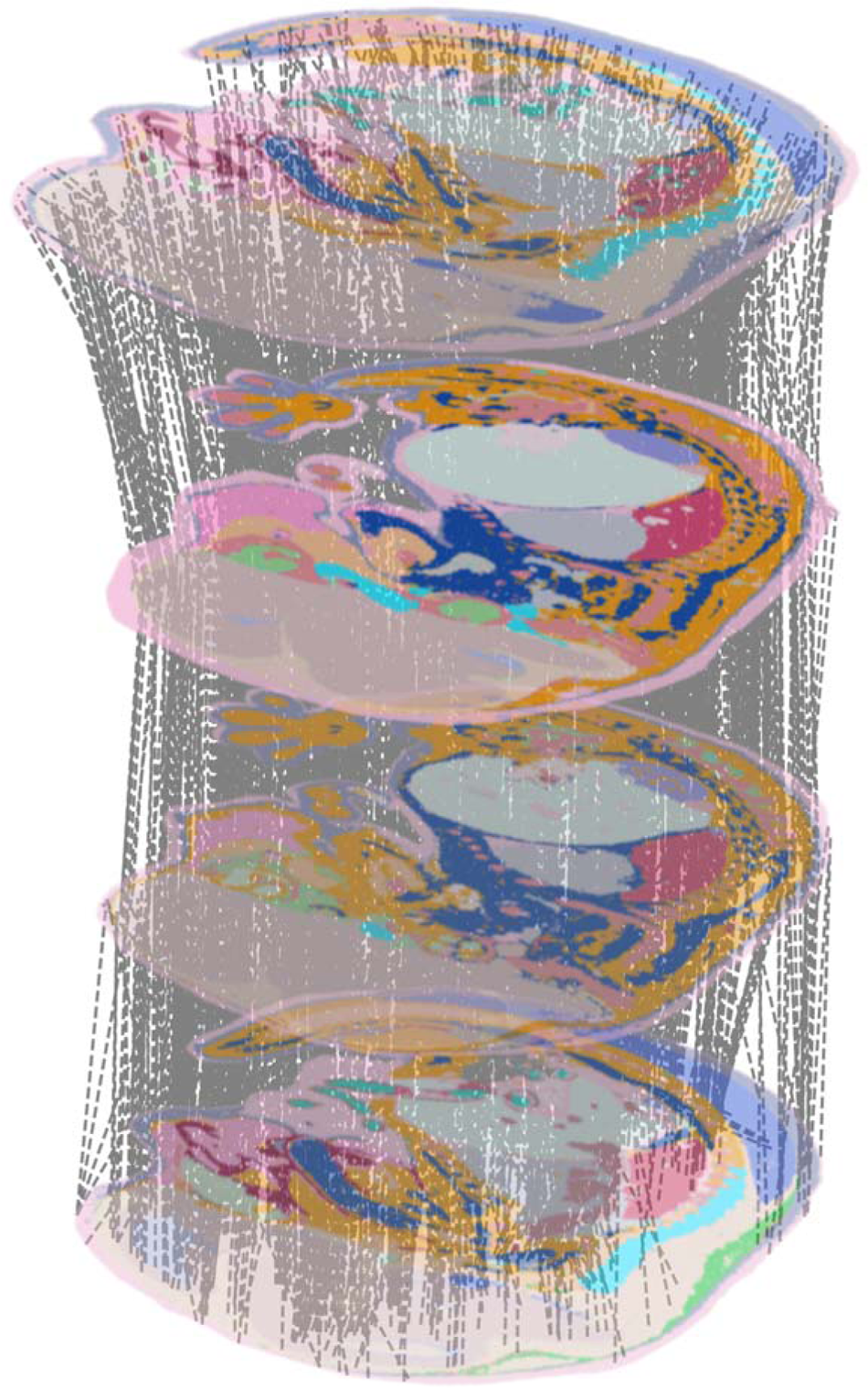
Multi-slice SLAT alignment for 3D reconstruction. Simultaneous SLAT alignment across four slices from the same E15.5 mouse embryo, ordered by vertical position (subsampled to 300 alignment pairs for clear visualization).

## Methods

### SLAT framework

#### Modeling spatial coordinates and omics feature of single cell

We denote a spatial omics dataset as *D = {(g^(i)^,s^(i)^),i = 1,2,…, N}*, where *N* is the number of spots or cells, g^(i)^ ∈ R^G^, and s^(i)^ ∈ R^2^ are the raw omics features (e.g., genes) and spatial coordinates of cell *i*, respectively, where *G* is the number of omics features. For datasets containing non-identical omics features, we use their overlapping features. For ease of notation, we denote the combination of omics features and spatial coordinates across all cells in a dataset as matrices G ∈ R^NxG^, S ∈ R^Nx2^, respectively. Subscripts such as G_1_,G_2_ and s_1_, s_2_ are added to distinguish two datasets being aligned.

In attempts to correct inter-sample batch effects, we employ an SVD-based cross-dataset matrix decomposition strategy as a preprocessing step. To begin with, we denote the log-normalized and scaled omics matrices of two spatial datasets as G_1_ ∈ R^N_1_xG^ and G_2_ ∈ R^N2xG^. We then apply SVD on their dot product as follows:

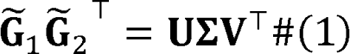

Using the decomposed matrices, we obtain the batch-corrected embeddings of the two datasets as:

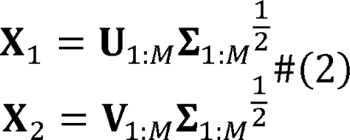

where U_1:M_ ∈ R^N^_1_^xM^, v_1:M_ ∈ R^N2xM^, E_1:M_ ∈ R^MxM^ are truncated decompositions corresponding to the *M*-largest singular values. Notably, this preprocessing step is flexible and allows for modular input (e.g., embedding produced by dedicated algorithms like GLUE^22^ for cross-modality integration).

We model the spatial information of cells in *D* as a spatial graph *G*= (v, e, x), where each node v_i_ ∈ *v* corresponds to a cell with the batch corrected embedding x^(i)^ ∈ R^M^ as its node attribute, and the edges connect the *K*-nearest neighbors in the spatial space. We also denote the adjacency matrix of the graph as A ∈ {0,1}^NxN^. A_i,j_ =1 if the edge (v_i_, v_j_) ∈ s, otherwise A_i,j_ = 0. In particular, for cross-technology alignment, different technologies may have distinct spatial resolutions, in which case we can select different *K*’s for each slice according to its spatial resolution. SLAT also supports building the spatial graph by radius, where all cells located within a specific radius are taken as “neighbors”.

SLAT formulates the alignment of two spatial datasets D_1_ = {(x_1_^(i)^,s_1_^(i)^),i = 1,2, …, N_1_} and D_2_ = {(x_2_^(i)^,s_2_^(i)^),i = 1,2, …, N_2_} as a minimum-cost bipartite matching problem of their corresponding spatial graphs G_1_ = (v_1_,s_1_,x_1_) and G_2_ = (v_2_,s_2_,x_2_):

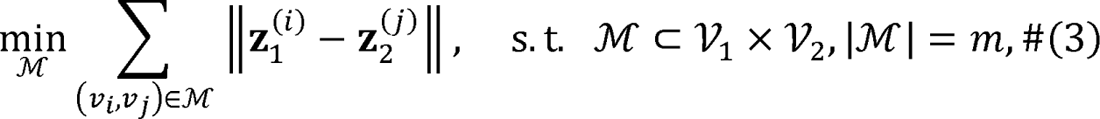

where Z_1_^(i)^, Z_2_^(j)^ ∈ R^P^ are node embeddings of v_i_ and v_j_ in G_1_ and G_2_, respectively, *M* is a set of matches of fixed size, and *P* is the dimensionality of node embeddings. Similarly, we use matrices z_1_ ∈ R^N^_1_^xP^ and z_2_ ∈ R^N2xP^ to denote the combination of cell embeddings of all cells in the two datasets.

It has been demonstrated in a previous work, that the above matching problem is equivalent to minimizing the Wasserstein distance between node embeddings from different graphs^16^. SLAT follows the same approach with adaptations for spatial omics data. Below, we explain how node or cell embeddings z_1_,z_2_ can be obtained and optimized for spatial graph alignment.

### Construction of holistic cell representations

An accurate alignment of spatial omics datasets should align cells that are similar in both the molecular modality and the spatial context. In particular, the spatial context can involve various resolutions, ranging from microenvironments to global positions within the tissue. Inspired by previous work^16, 36, 37^, we first employ the lightweight graph-convolutional network (LGCN) to derive a holistic cell representation with all such information integrated for each dataset. A LGCN propagates and aggregates information along the spatial graph through stepwise concatenations:

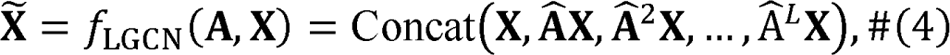

where 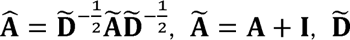
 is the diagonal degree matrix of A^∼^, and *L* is the maximal number of steps. The resulting cell representation x^∼^ ∈ R^Nx(L+1)M^ is a concatenation of multi-level information. The first *M* dimensions correspond to no graph propagation, which is simply a copy of the omics data X. The second *M* dimensions correspond to one-step graph propagation, reflecting the composition of a cell’s immediate neighbors, which form its microenvironment. The information coarsens as the number of steps increases, gradually becoming a representation of rough locations within the tissue. Thus, x∼ contains informative features for spatial alignment at multiple levels of spatial context.

The cell representation x∼ is constructed separately for each dataset by using dataset-specific omics profiles and adjacency matrices:

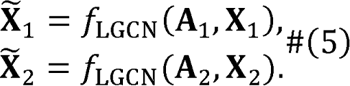

### Adversarial graph alignment

Based on the holistic cell representations x^∼^_1_, x^∼^_2_ described above, we use adversarial alignment to learn cell embeddings z_1_,z_2_ that minimize the Wasserstein distance for graph matching^16^. Specifically, we apply a multilayer perceptron denoted as *f_Z_* to mitigate systematic bias in x∼_1_ and x∼_2_ that may arise from differences in omics distribution or spatial topology across datasets:

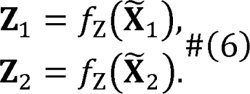

We then introduce the Wasserstein discriminator f_D_, which uses z_1_, z_2_ as input and tries to maximize the following Wasserstein loss L_W_ to estimate the Wasserstein distance:

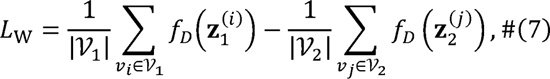

where Z_1_^(i)^ is a row in z_1_ corresponding to cell *v*_i_, and Z_2_^(j)^ is a row in z_2_ corresponding to cell v_j_. The transformation f_Z_ can then be adversarially trained to minimize (7), for aligning the distribution of cell embeddings in the two datasets properly. However, different single-cell spatial datasets may contain different cell-type proportions or distinct spatial regions. It is thus unreasonable to assume identical distribution of their cell embeddings as assumed in the standard scheme described above^38^. Inspired by a previous study^16^, we use the output of the Wasserstein discriminator *f*_D_ as a dynamic clipping criterium to select lx N_1_ and lx N_2_ cells from the two datasets with minimum Wasserstein distance for adversarial training (Extended Data Fig. 1a):

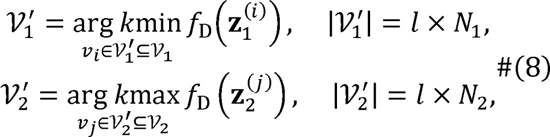

where *l* is a hyperparameter between 0 and 1. These cells correspond to the most reliable anchors to guide the alignment. The Wasserstein discriminator loss L_W_ is then modified accordingly as follows:

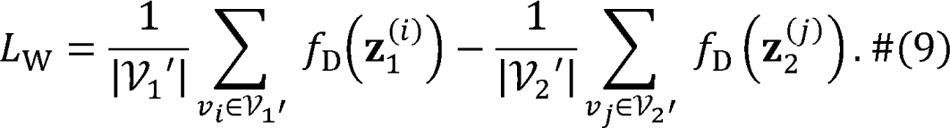

This approach ensures that distinct regions across two spatial datasets will not be forcibly aligned. The results show that SLAT performs best when *l* = 0.6, although the performance depends only weakly on *l* (Extended Data Fig. 2a).

To avoid a degenerate solution where all embeddings collapse to a singular point, we adopt an additional reconstruction term to ensure that the embeddings have sufficient information to reconstruct input, essentially enhancing model stability. We use a simple multilayer perceptron network denoted as *f*_R_ for data reconstruction, making the following reconstruction loss:

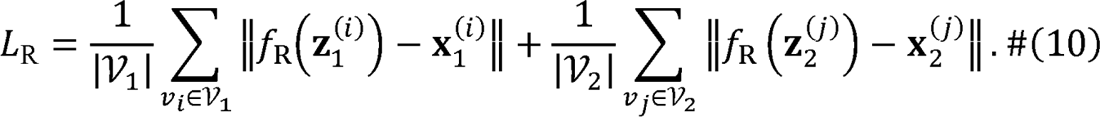

### Overall objective

Finally, the overall training objective of SLAT can be summarized as follows:

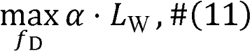

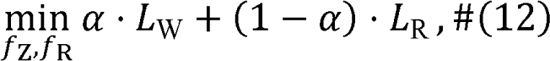

where *L*_W_ and *L*_R_ are defined by Eqs. (9) and (10), respectively, a is a hyperparameter balancing the contribution of adversarial alignment and data reconstruction. We use stochastic gradient descent (SGD) with the Adam optimizer to train the SLAT model.

### Coordinate matching

Apart from the core model described above, we also provide options to use additional information to match the spatial coordinates **s** of two slices which can help distinguish symmetric structures (e.g., left and right hemispheres of the brain) and improve the final matching quality. The goal of coordinate matching is to roughly align different slices in terms of their overall direction by estimating an affine transformation matrix **M**. The exact strategy depends on the type of information available.

First, guided by expert knowledge of tissue structures, **M** can be computed by combining the scaling, rotation, and translation operations required to obtain a rough alignment between the two slices. Second, if imaging data like H-E staining images are available, **M** can be estimated following our tutorial based on SimpleElastix^39^, which is a state-of-the-art medical image registration tool. Finally, if no other information is available, we also provide a default solution based on iterative closet point (ICP)^40^, a point-cloud registration algorithm, where we treat the spatial datasets as point clouds on a two-dimensional plane and uses geometric features for registration. With the obtained **M** matrix, we consort the coordinates of the two datasets by the following transformation:

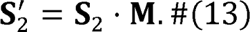

### Cell matching and quality assessment

With the cell embeddings z_1_, z_2_ learned by the SLAT core model and the matched spatial coordinates s_1_ and s’_2_, we match dataset *D*_2_ with *D*_1_ using the following strategy: For cell *j* in dataset *D*_2_, SLAT first selects the top *K* most similar *D*_1_ cells in the embedding space z as a candidate set c_j_, then chooses the one with minimal distance in matched spatial coordinates s as the final match.

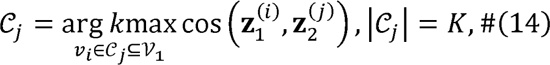

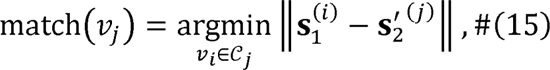

where s_1_^(i)^ and s_2_^’ (j)^ are rows in the coordinate matrices s_1_ and s_2_^’^, respectively. *K* is a hyperparameter that can be adjusted according to the confidence in the coordinate matching result. The cosine similarity cos (Z_1_^(i)^, Z_2_^(j)^) is used to quantify the matching quality. We filter out unreliable matches by defining a cutoff for cosine similarity. Cell pairs with low matching quality suggest either biologically distinct changes or technological artifacts and should be carefully checked (see Results). Empirically, we introduce a default cutoff at 0.8 which can also be adjusted in a case-by-case manner.

### Systematic benchmarks

#### Benchmark datasets

We selected 10x Visium, MERFISH and Stereo-seq as representative spatial technologies for benchmarking alignment methods. The 10x Visium dataset comes from consecutive slices of human dorsolateral prefrontal cortex, containing about 3,000 spots per slice, each spanning 50 μm with over 20,000 genes detected^41^. The MERFISH dataset comes from consecutive slices of mouse hypothalamic preoptic, with subcellular resolution but only 151 genes detected in total^21^. The Stereo-seq dataset comes from consecutive slices of an E15.5 mouse embryo, containing over 100,000 single cells per slice, divided into 25 cell types with complex spatial organization, and detects over 20,000 genes in total^8^. For each technology, we chose the first two slices if the dataset contains more than two slices. Cell type annotation and tissue region segmentation were obtained from the original authors whenever possible. Since the Stereo-seq dataset does not provide tissue segmentation, we segmented the most prominent regions in the embryo, including Brain, Jaw and face, Spinal cord, Heart, Lung, Liver, and Belly under the guidance of an expert in mouse anatomy. The spatial segmentation of the MERFISH dataset is provided in publication figures but not in raw data, so we re-segmented the data as guided by the figures.

#### Slice processing

For each technology, we first removed the cells/spots that are unannotated in both slices, then rotated the second slice with a random angle before feeding to the alignment methods.

#### Benchmarked methods

The benchmarked methods Harmony, PASTE, STAGATE, and SLAT were executed using the Python packages “harmonypy” (v0.0.6), “paste-bio” (v1.3.0), “STAGATE_pyG” (latest commit 8b9c8ef), and “scSLAT” (v0.2.0), respectively, in Python (v3.8). Seurat was executed using the R package “Seurat” (v4.1.1) in R (v4.1.3). For each method, we used the default hyperparameter settings and data preprocessing steps recommended by the original authors.

#### Evaluation metrics

Intuitively, a proper spatial alignment should align cells matched in both molecular profile and spatial context. Thus, we reported the cell type matching accuracy and spatial region matching accuracy simultaneously via the form of confusion matrices. For convenience, we also defined a “joint accuracy” as the proportion of cells with both cell type and spatial region matched correctly, which corresponds to the upper right corner of the confusion matrix.

Meanwhile, we also noticed that there exists other metrics commonly used for evaluating graph alignment, e.g., the edge score^42, 43^. However, we did not use it as we found that it can be deceived in certain situations. For example, Extended Data Fig. 1b shows two graphs with known ground truth node pairing information. “Alignment 1” is the correct matching (grey lines) and “Alignment 2” has four mismatched pairs (highlighted by red lines), but the two alignments get the same edge score.

#### Benchmark tasks

We benchmarked the alignment methods based on the following four tasks: (1) real world alignment, (2) split slice alignment, (3) scalability test:

In real world alignment, the slices to be aligned are different slices produced from the same position of the same tissue^5, 8, 41^ (Supplementary Fig. 1). Each algorithm was run eight times with different model random seeds. Performance of the alignment methods is presented as the confusion matrix as described above.

In split slice alignment, we split the first slice of each technology used in real world alignment into two pseudo-slices of equal size by randomly sampling the cells without replacement (Supplementary Fig. 3). Each algorithm was run eight times with different model random seeds. Performance quantification is the same as in real world alignment.

For scalability benchmark, we randomly subsampled the Stereo-seq dataset used in real world benchmark to a range of cell numbers (3200, 6400, 12800, 25600, 51200, 102400). The subsampling process was repeated eight times with different random seeds (Supplementary Fig. 5).

#### Benchmark workflow

We used Snakemake (v7.12.0) to manage the whole benchmark workflow. All benchmarked methods were allocated 16 cores of Intel Xeon Platinum 8358 CPU, 128 GB of RAM, and a NVIDIA A100 GPU with 80 GB VRAM by the Slurm workload management system.

#### Hyperparameter robustness

We tested SLAT’s robustness to key hyperparameters including: (1) neighbor size *K* for the spatial graph, (2) input embedding methods, (3) number of training epochs, (4) number of LGCN layers *L*, (5) MLP hidden layer dimension, (6) dimension *P* of SLAT embedding, (7) weight a of the adversarial loss, (8) dynamic clipping ratio l, (9) learning rate of the SLAT model.

We run SLAT in two homogeneous Stereo-seq datasets from the same E15.5 mouse embryo, each containing more than 100,000 cells so we randomly subsampled 8,000 cells from each dataset to save time. Every experiment was run 8 times with different model random seeds (Extended Data Fig. 2).

#### Robustness to noise in the spatial graph

In practice, the spatial graph may be imperfect due to technical limitations. Therefore, we tested SLAT’s robustness to graph corruption in the same dataset used in hyperparameter robustness evaluation. Specifically, we randomly masked the edges in the graph by increasing ratios (from 0.1 and 0.9). Every experiment was run 8 times with different masking random seeds.

### Heterogeneous alignment across distinct technologies and modalities

#### Visium and Xenium data alignment

The Visium and Xenium datasets were generated from consecutive slices of the human breast cancer tissue sample^4^. In order to maintain the comparability with the original paper, we chose the exact same slices used in their analysis (see Fig. 4c of ref^4^). We further selected the shared region between the two slices as the original authors reported^4^ for follow-up analysis.

Considering that the Xenium slice contains more than 100,642 cells while Visium only contains less than 3,841 spots in same physical region, we used different neighbor sizes proportional to cell density (*K*= 5 for Visium, and *K*= 130 for Xenium) when constructing the spatial graphs, in order to ensure that the GCNs have similar spatial receptive fields. SLAT is then run with otherwise default parameters. We selected Visium triple positive spots based on the number of aligned Xenium triple positive cells (Supplementary Fig. 6b). 7 spots with more than 2 aligned cells were chosen. Following the original paper, we did differential gene expression analysis of the SLAT identified and manually curated triple positive spots against all other spots, respectively, using the Scanpy^44^ function “scanpy.tl.rank_genes_groups” with parameter “methods=‘wilcoxon’”. For comparison, all methods included in the benchmarks (PASTE, Harmony, Seurat, and Harmony) were applied to the same Visium and Xenium datasets with their default parameters.

For comparison, all methods benchmarked (PASTE, Harmony, Seurat, and Harmony) were applied to the same datasets with their default parameters (Extended Data Fig. 5).

#### SeqFISH data and Stereo-seq data alignment

For Stereo-seq and seqFISH, we chose the E9.5 slice with the most complete cell type annotation by the original authors (i.e., the slices with the least proportion of unannotated cells), respectively. To align the chosen seqFISH and Stereo-seq slices, we run SLAT using the default parameters and filtered out alignment pairs with similarity scores lower than 0.8. We next refined cell type annotations in the Stereo-seq dataset based on the higher resolution annotation in seqFISH through label transfer for “Neural crest” cells (Fig. 3b). We also manually annotated “Neural crest” cells for independent validation. Scanpy^44^ was used for the analysis following its official tutorial: the data were log-normalized using the functions “sc.pp.normalize_total” and “sc.pp.log1p”. Highly variable genes were identified with “sc.pp.highly_variable_genes”. The first 50 principal components after PCA (“sc.tl.pca”) were used to generate neighborhood graphs (“sc.pp.neighbors”) for computing UMAP embeddings (“sc.tl.umap”) and Leiden clustering (“sc.tl.leiden”). The “Neural crest” cells were annotated via marker genes of different germ layers (*Foxc2* and *Vcan* for mesoderm, *Msx1*, *Mif*, and *Dik1*for ectoderm, see Extended Data Fig. 3d).

For comparison, all methods benchmarked (PASTE, STAGATE, Seurat and Harmony) were applied to the same datasets with their default parameters (Extended Data Fig. 4).

#### Spatial-ATAC-seq and Stereo-seq data alignment

For cross-modality alignment, the slices to align were two 11.5-day mouse embryo datasets from Stereo-seq (RNA) and spatial-ATAC-seq (ATAC), respectively. For Stereo-seq, we chose the E11.5 slice with the most complete cell type annotation by the original authors (i.e., the slice with the least proportion of unannotated cells). For spatial-ATAC-seq, we chose the E11.5 slice with the highest spatial resolution (20 μm). Given that the current spatial-ATAC-seq data did not cover the entire embryo due to technical limitations, we extracted the anatomically corresponding regions from the Stereo-seq dataset under expert guidance.

To project cells from different modalities into a shared latent space, we employed the graph-linked multi-modality embedding strategy we proposed before ^22^, then ran SLAT with the default hyperparameters. Based on the outputted alignment, we transferred cell type labels from Stereo-seq to spatial-ATAC-seq which was not annotated, and then applied SCENIC+^45^ for joint regulatory inference.

To compare with benchmarked methods (PASTE, Harmony, Seurat and Harmony), we also used the same multi-modality embedding as their input followed by their default pipeline. Exceptions are Seurat and STAGATE, which do not support low dimensional embeddings as input, thus cannot be compared. In addition, we also compared with the original multi-modality embeddings^22^ directly (Supplementary Fig. 7).

#### Spatial-temporal alignment

We applied SLAT to align E11.5 and E12.5 heterogeneous mouse embryo Stereo-seq datasets with the default hyperparameters. In order to maintain the comparability with the original paper, we chose the exact same slices used in their analysis (see Fig. 3a of ref^8^). Regions with lower SLAT similarity scores were marked in Fig. 5b. To focus on kidney development, we extracted cells labeled as “Kidney” in E12.5 and their aligned cells in E11.5, then clustered these cells by using the standard Scanpy clustering pipeline mentioned above and annotated them via well-defined kidney markers^31, 32^ (*Osr1*, *Foxc1* and *Podxl* for nephron progenitors; *Uncx*, *Nr2f2*, *Dach*, *Wt1*, *Nphs1*, and *Cd44* for kidney; see Fig. 4f). To further demonstrate the robustness of SLAT, we rerun the analysis against two additional slices randomly chosen from E11.5 and E12.5, and obtained similar results (Supplementary Fig. 8).

For comparison, all methods benchmarked (PASTE, STAGATE, Seurat and Harmony) were applied to the same datasets with their default parameters (Extended Data Fig. 8).

## Data availability

All datasets used in this study were already published and were obtained from public data repositories. See Supplementary Table 1 for detailed information on the entire datasets used in this study, including publication and downloading URLs.

## Code availability

The SLAT framework was implemented in the “scSLAT” Python package, which is available at https://github.com/gao-lab/SLAT. For reproducibility, the scripts for all benchmarks were assembled using Snakemake (v7.12.0), which is also available in the above repository.

