## Supplementary-Figure for "Spatial-linked alignment tool (SLAT) for aligning heterogenous slices properly"

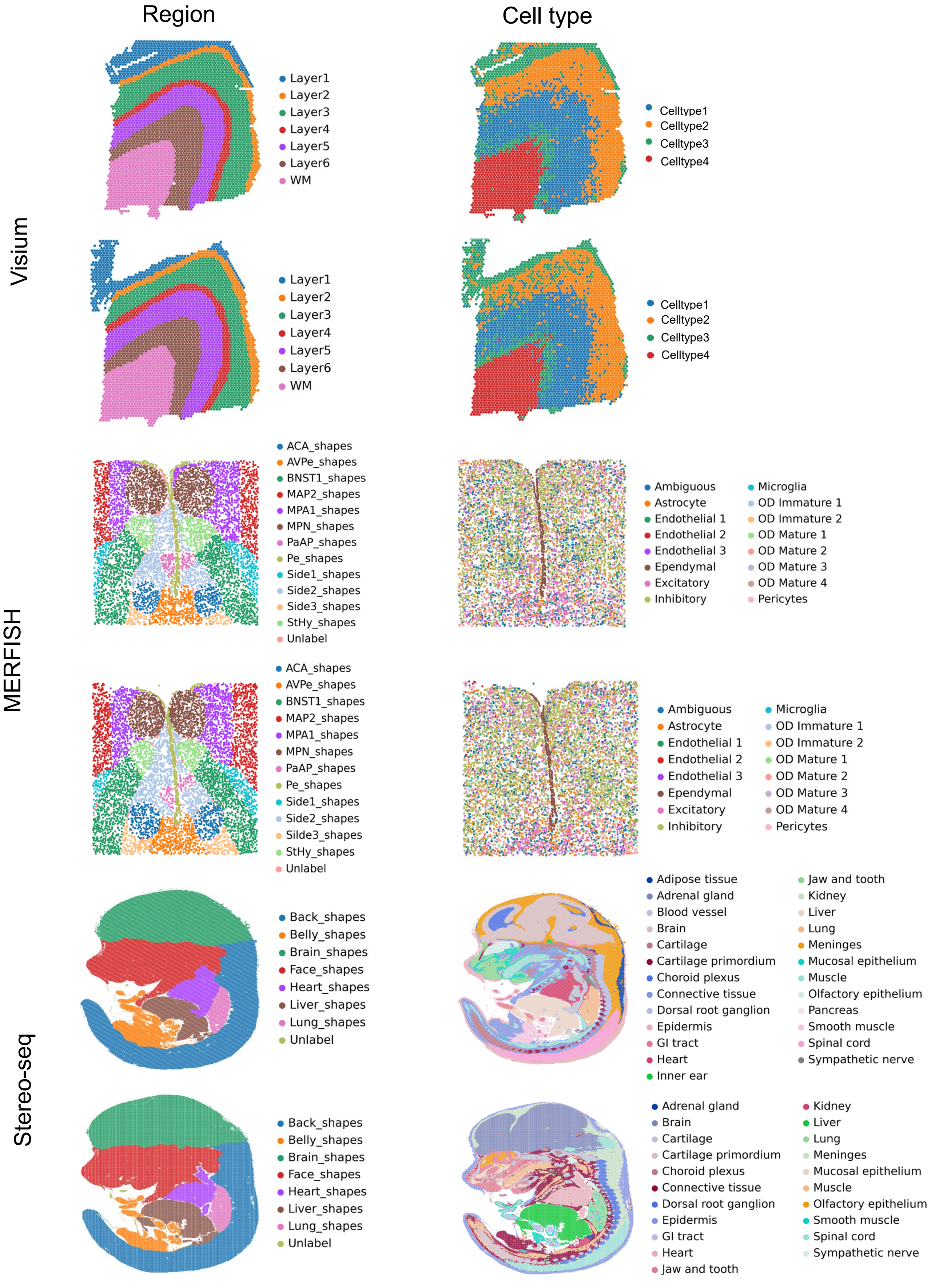


**Supplementary Fig. 1 Visualization of benchmark datasets.**
Visualization of slices used in the benchmarks. slices are colored by spatial regions (left panels) and cell types (right panels), respectively

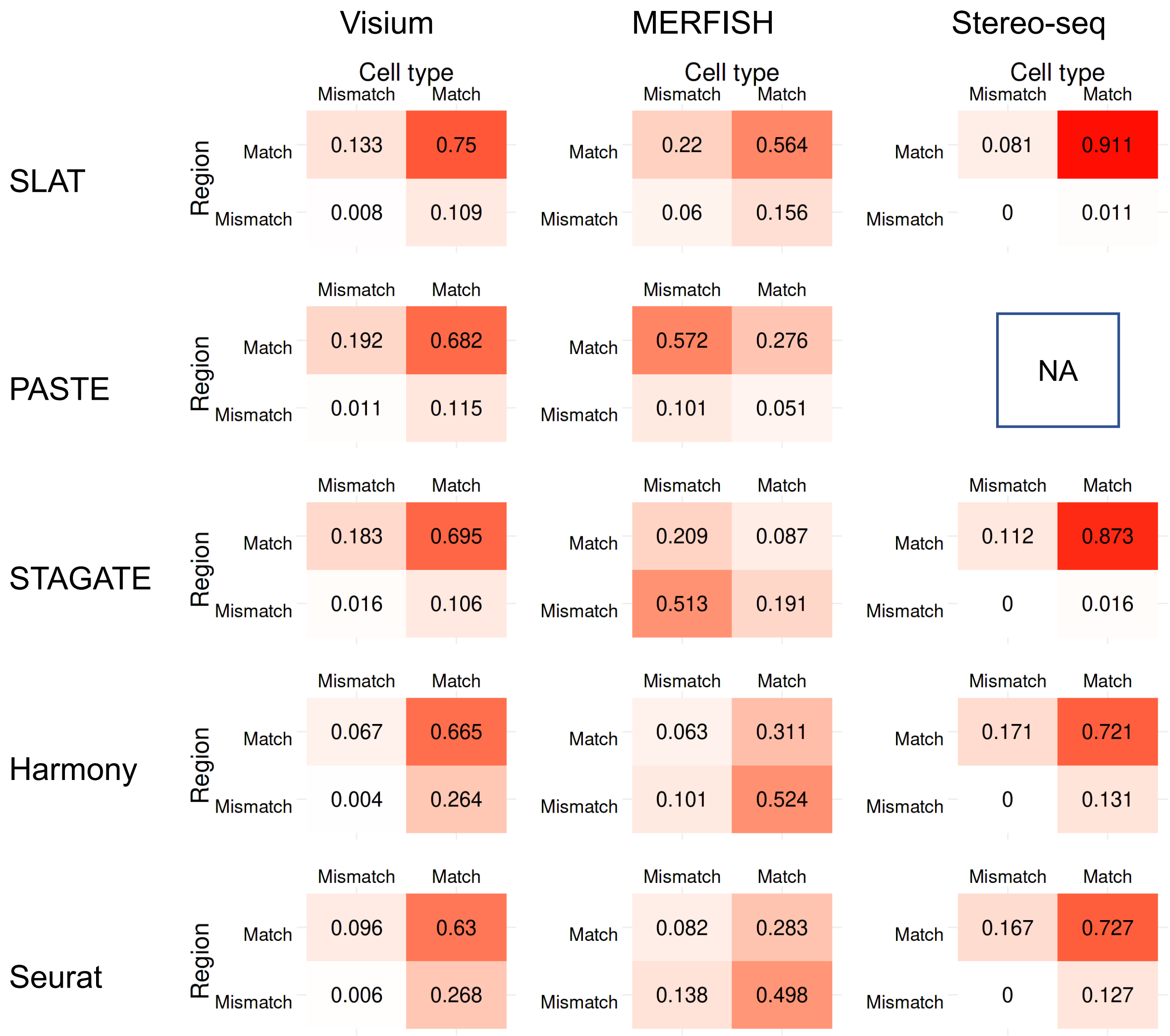


**Supplementary Fig. 2 Evaluation different methods on split spatial datasets.**
Heatmaps quantifying the region matching accuracy and cell type matching accuracy of SLAT, PASTE, STAGATE, Harmony and Seurat respectively in the form of confusion matrices of split dataset in Supplementary Fig. 3. The number in each cell is the average proportion across eight repeats with different random seeds. PASTE failed to run on the Stereo-seq dataset due to GPU memory overflow (capping at 80 GB).

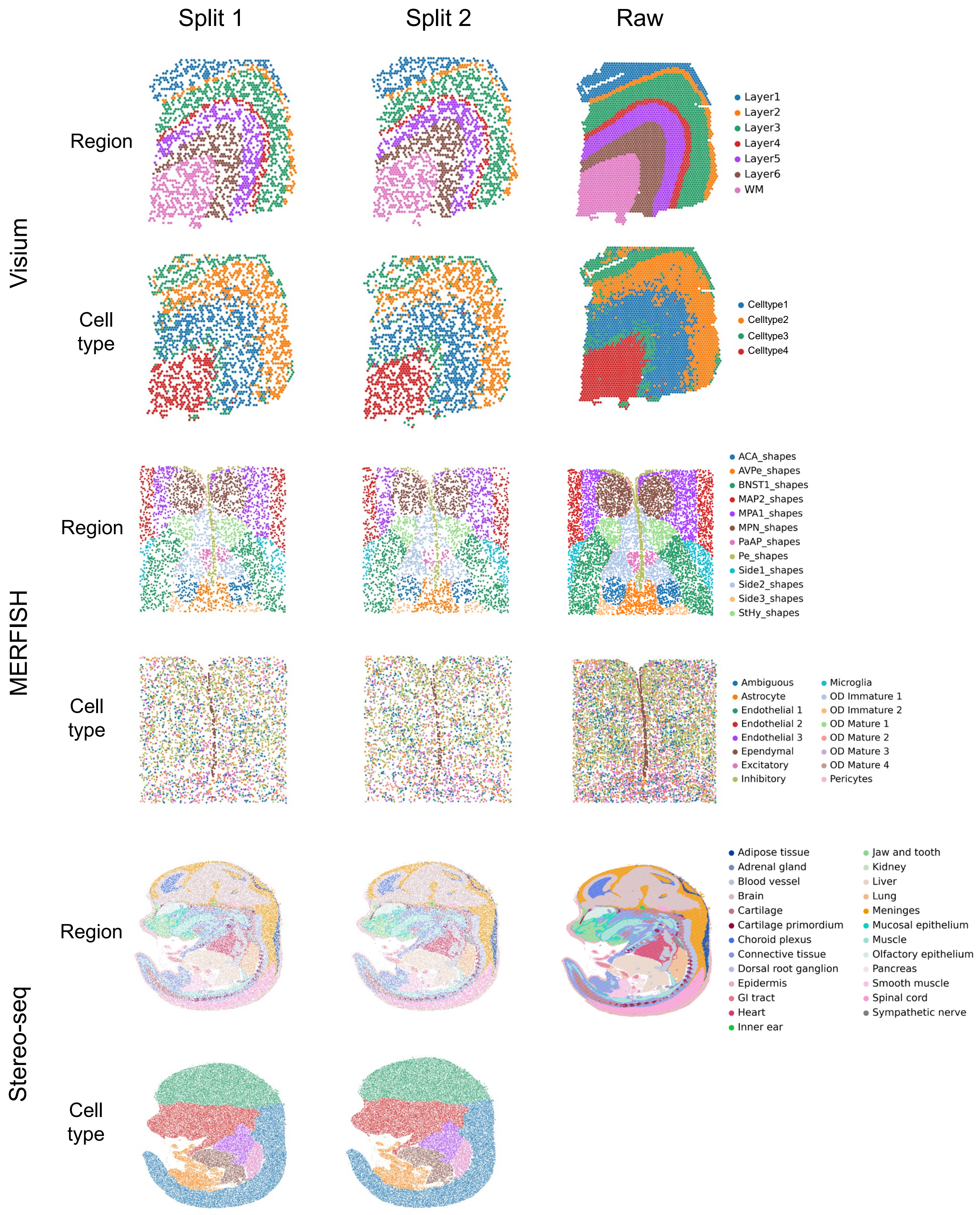


**Supplementary Fig. 3 Visualization of split datasets.**
Visualization of randomly split slices and raw slices used in the benchmark. Slices are colored by spatial regions and cell types, respectively.

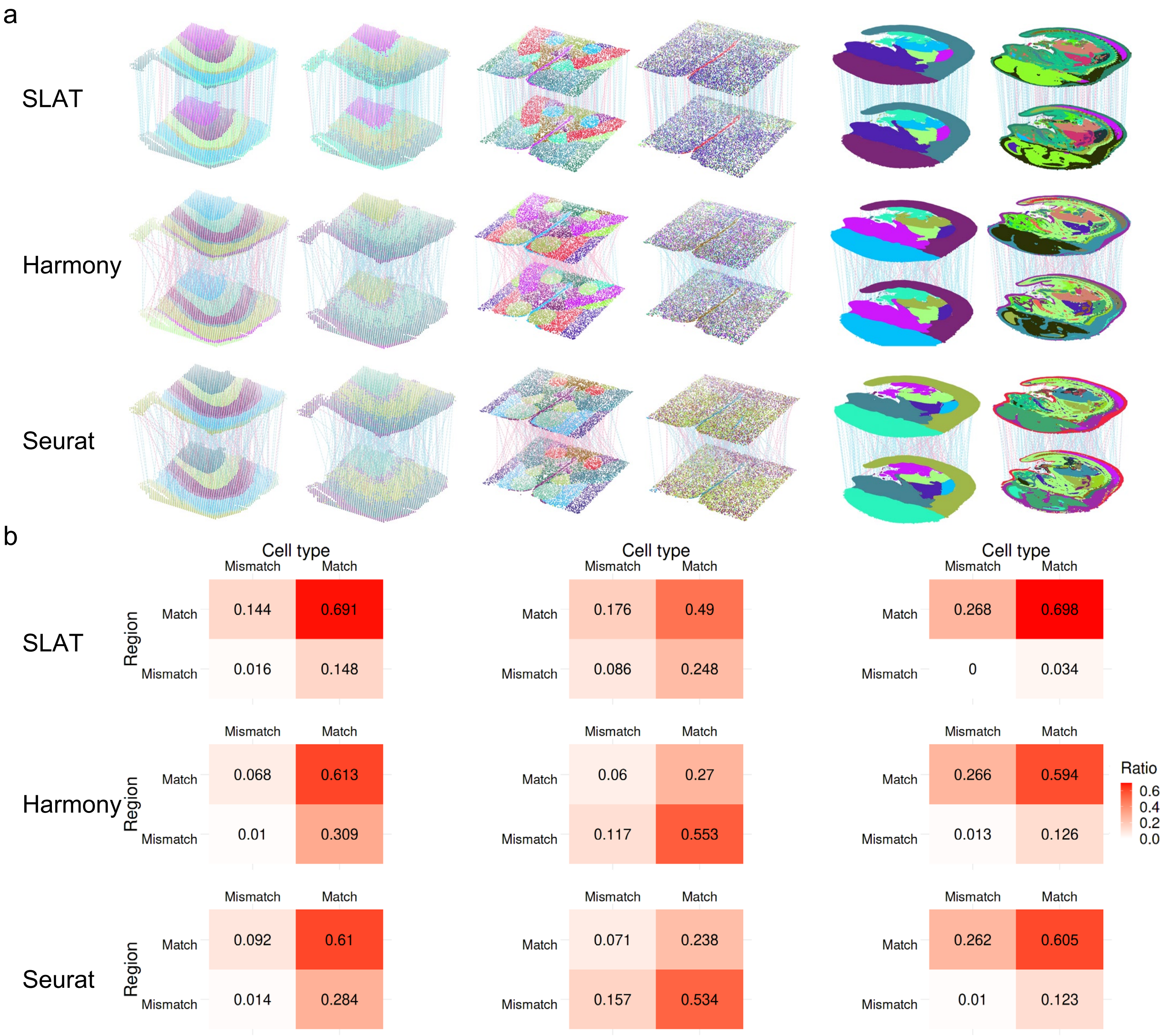


**Supplementary Fig. 4 Visualization of evaluation SLAT on homogeneous spatial alignment with spatially unaware algorithms.**

**a**, Visualization of alignment results of different methods on the benchmark datasets in Fig. 2a. Vertical lines connect aligned cell pairs (subsampled to 300 alignment pairs for clear visualization). Blue lines indicate correct alignments (with matching cell types and spatial regions) while red lines indicate incorrect alignments. **b**, Heatmaps quantifying the region matching accuracy and cell type matching accuracy respectively in the form of confusion matrices for the alignments shown in **a**.

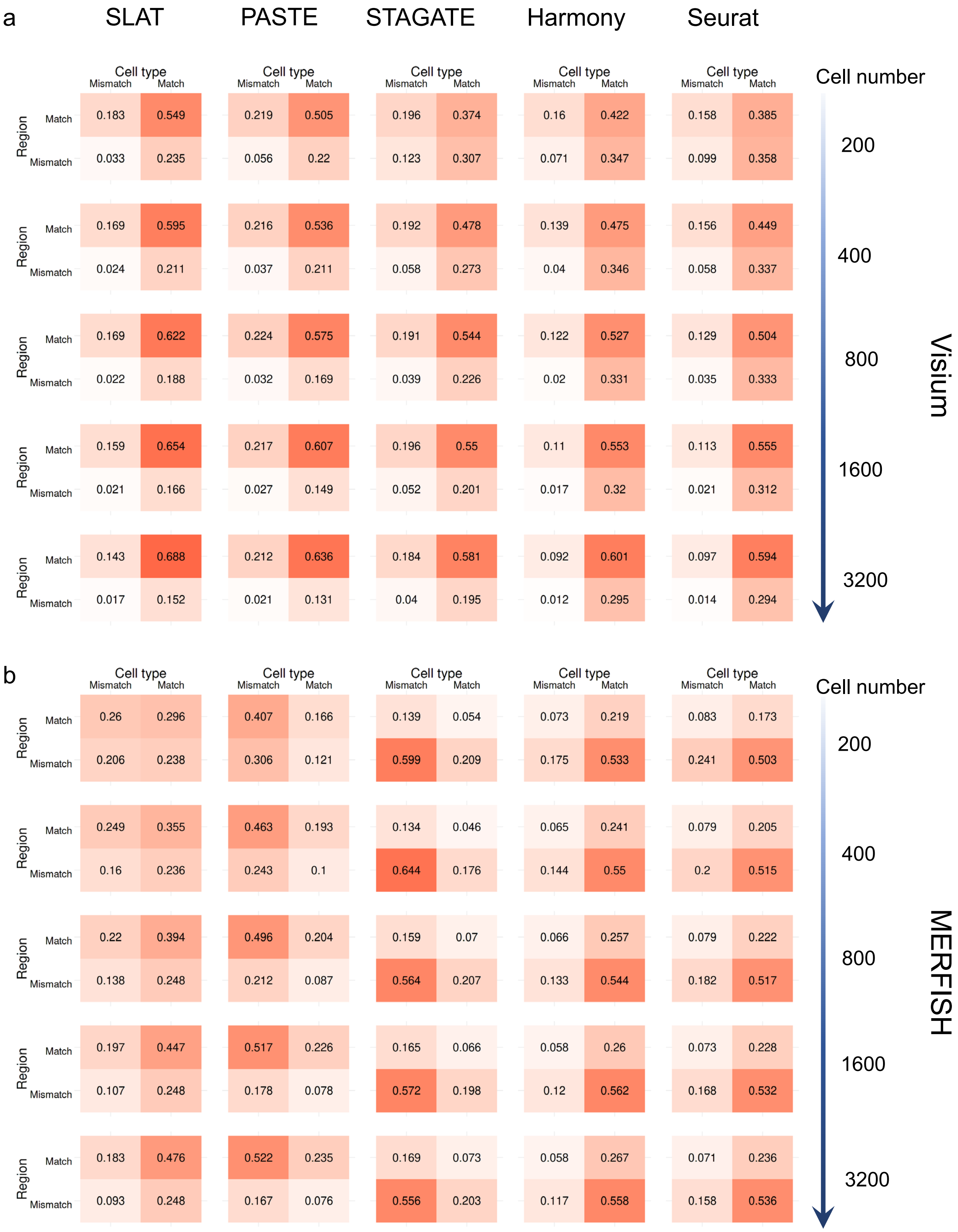

a
SLAT
PASTE
STAGATE
Harmony
Seurat
Cell type
Mismatch
Match
Region
Match
Mismatch
Cell number
200
400
800
1600
3200
Visium
0.183 0.549 0.033 0.235
0.219 0.505 0.056 0.22
0.196 0.374 0.123 0.307
0.16 0.422 0.071 0.347
0.158 0.385 0.099 0.358
0.169 0.595 0.024 0.211
0.216 0.536 0.037 0.211
0.192 0.478 0.058 0.273
0.139 0.475 0.04 0.346
0.156 0.449 0.058 0.337
0.169 0.622 0.022 0.188
0.224 0.575 0.032 0.169
0.191 0.544 0.039 0.226
0.122 0.527 0.02 0.331
0.129 0.504 0.035 0.333
0.159 0.654 0.021 0.166
0.217 0.607 0.027 0.149
0.196 0.55 0.052 0.201
0.11 0.553 0.017 0.32
0.113 0.555 0.021 0.312
0.143 0.688 0.017 0.152
0.212 0.636 0.021 0.131
0.184 0.581 0.04 0.195
0.092 0.601 0.012 0.295
0.097 0.594 0.014 0.294
b
MERFISH
0.26 0.296 0.206 0.238
0.407 0.166 0.306 0.121
0.139 0.054 0.599 0.209
0.073 0.219 0.175 0.533
0.083 0.173 0.241 0.503
0.249 0.355 0.16 0.236
0.463 0.193 0.243 0.1
0.134 0.046 0.644 0.176
0.065 0.241 0.144 0.55
0.079 0.205 0.2 0.515
0.22 0.394 0.138 0.248
0.496 0.204 0.212 0.087
0.159 0.07 0.564 0.207
0.066 0.257 0.133 0.544
0.079 0.222 0.182 0.517
0.197 0.447 0.107 0.248
0.517 0.226 0.178 0.078
0.165 0.066 0.572 0.198
0.058 0.26 0.12 0.562
0.073 0.228 0.168 0.532
0.183 0.476 0.093 0.248
0.522 0.235 0.167 0.076
0.169 0.073 0.556 0.203
0.058 0.267 0.117 0.558
0.071 0.236 0.158 0.536

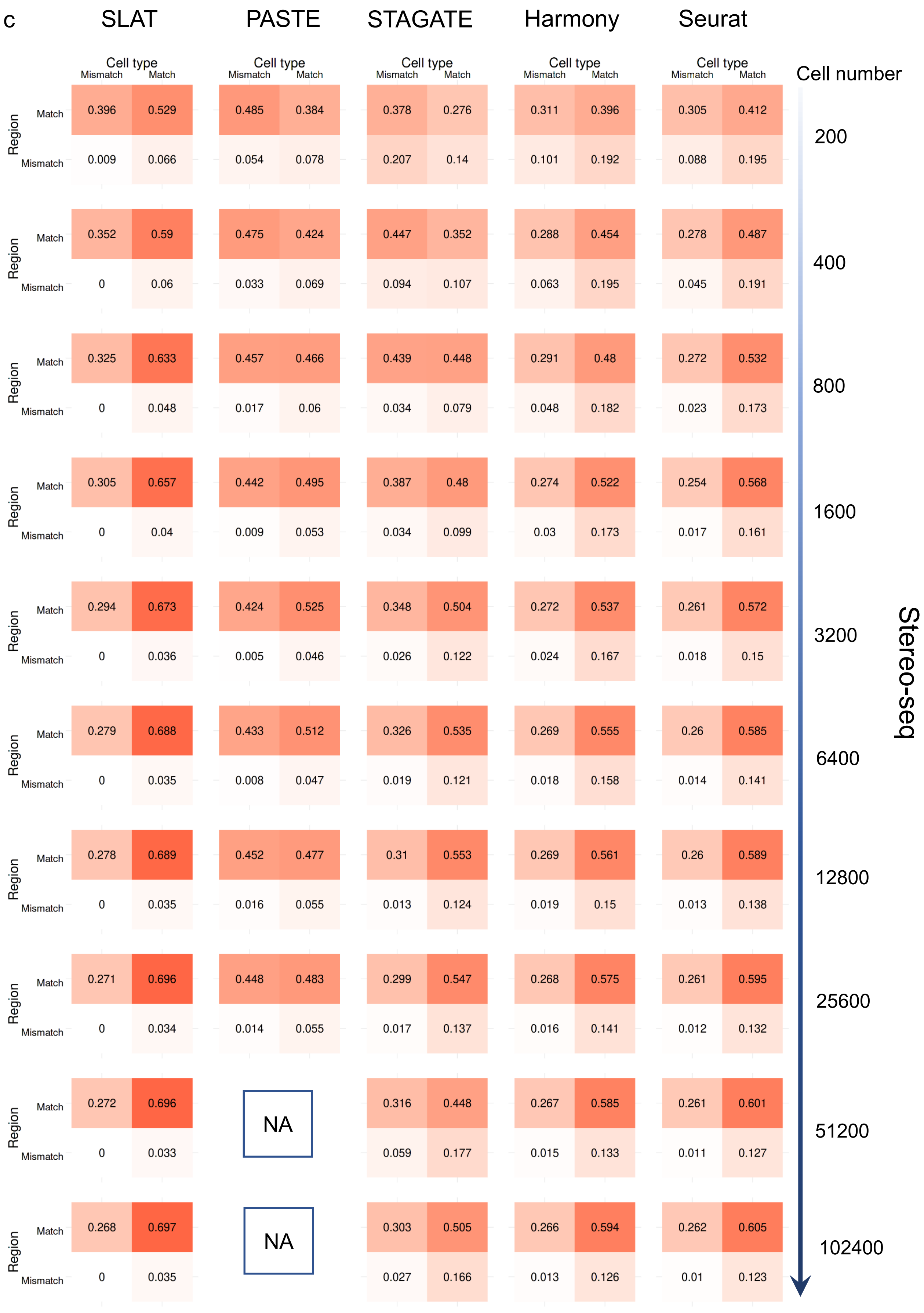
c
SLAT
PASTE
STAGATE
Harmony
Seurat
Cell type
Mismatch
Match
Region
Match
Mismatch
Cell number
Stereo-seq
200
0.396 0.529 0.485 0.384 0.378 0.276 0.311 0.396 0.305 0.412
0.009 0.066 0.054 0.078 0.207 0.14 0.101 0.192 0.088 0.195
400
0.352 0.59 0.475 0.424 0.447 0.352 0.288 0.454 0.278 0.487
0 0.06 0.033 0.069 0.094 0.107 0.063 0.195 0.045 0.191
800
0.325 0.633 0.457 0.466 0.439 0.448 0.291 0.48 0.272 0.532
0 0.048 0.017 0.06 0.034 0.079 0.048 0.182 0.023 0.173
1600
0.305 0.657 0.442 0.495 0.387 0.48 0.274 0.522 0.254 0.568
0 0.04 0.009 0.053 0.034 0.099 0.03 0.173 0.017 0.161
3200
0.294 0.673 0.424 0.525 0.348 0.504 0.272 0.537 0.261 0.572
0 0.036 0.005 0.046 0.026 0.122 0.024 0.167 0.018 0.15
6400
0.279 0.688 0.433 0.512 0.326 0.535 0.269 0.555 0.26 0.585
0 0.035 0.008 0.047 0.019 0.121 0.018 0.158 0.014 0.141
12800
0.278 0.689 0.452 0.477 0.31 0.553 0.269 0.561 0.26 0.589
0 0.035 0.016 0.055 0.013 0.124 0.019 0.15 0.013 0.138
25600
0.271 0.696 0.448 0.483 0.299 0.547 0.268 0.575 0.261 0.595
0 0.034 0.014 0.055 0.017 0.137 0.016 0.141 0.012 0.132
51200
0.272 0.696 NA 0.316 0.448 0.267 0.585 0.261 0.601
0 0.033 0.059 0.177 0.015 0.133 0.011 0.127
102400
0.268 0.697 NA 0.303 0.505 0.266 0.594 0.262 0.605
0 0.035 0.027 0.166 0.013 0.126 0.01 0.123

**Supplementary Fig. 5 Evaluation different methods on subsampled datasets.**
Heatmaps quantifying the performance changes of SLAT, PASTE, STAGATE, Harmony and Seurat with subsamples of varying sizes (labeled in figure) on Visium (**a**), MERFISH (**b**) and Stereo-seq (**c**) datasets, respectively. The number in each cell is the average proportion across eight repeats with different random seeds. PASTE failed to run on the Stereo-seq dataset due to GPU memory overflow (capping at 80 GB).

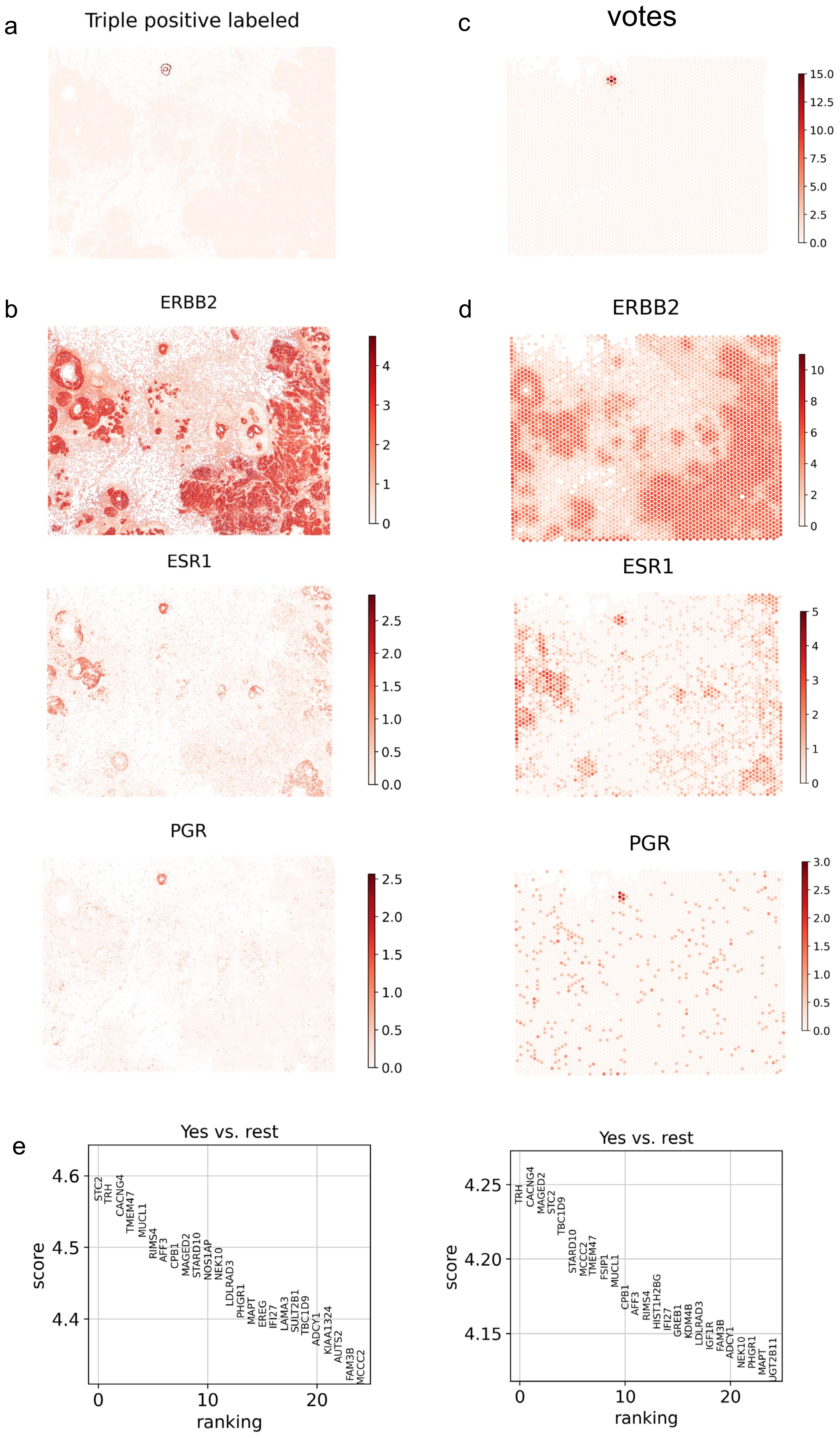
a
Triple positive labeled
c
votes
b
ERBB2
ESR1
PGR
d
ERBB2
ESR1
PGR
e
Yes vs. rest
score
ranking
STC2 TRH CACNG4 TMEM47 MUCL1 RIMS4 AFF3 CPB1 MAGED2 STARD10 NOS1AP NEK10 LDLRAD3 PHGR1 MAPT EREG IFI27 LAMA3 SULT2B1 TBC1D9 ADCY1 KIAA1324 AUTS2 FAM3B MCCC2
Yes vs. rest
score
ranking
TRH CACNG4 MAGED2 STC2 TBC1D9 STARD10 MCCC2 TMEM47 FSIP1 MUCL1 CPB1 AFF3 RIMS4 HIST1H2BG IFI27 GREB1 KDM4B LDLRAD3 IGF1R FAM3B ADCY1 NEK10 PHGR1 MAPT

### Supplementary Fig. 6 Xenium and Visium datasets.

**a**, Manually annotated triple positive breast tumor cells in the Xenium slice. **b**, **d**, Expression of three markers of triple positive breast cancer (*ERBB2*, *ESR1*, *PGR*) in the Visium (**b**) and Xenium (**d**) dataset. **c**, Number of Xenium cells aligned to each Visium spot. **f, g**, Top 25 highly expressed genes of the SLAT aligned (**f**) and manual curated (**g**) triple positive spots.

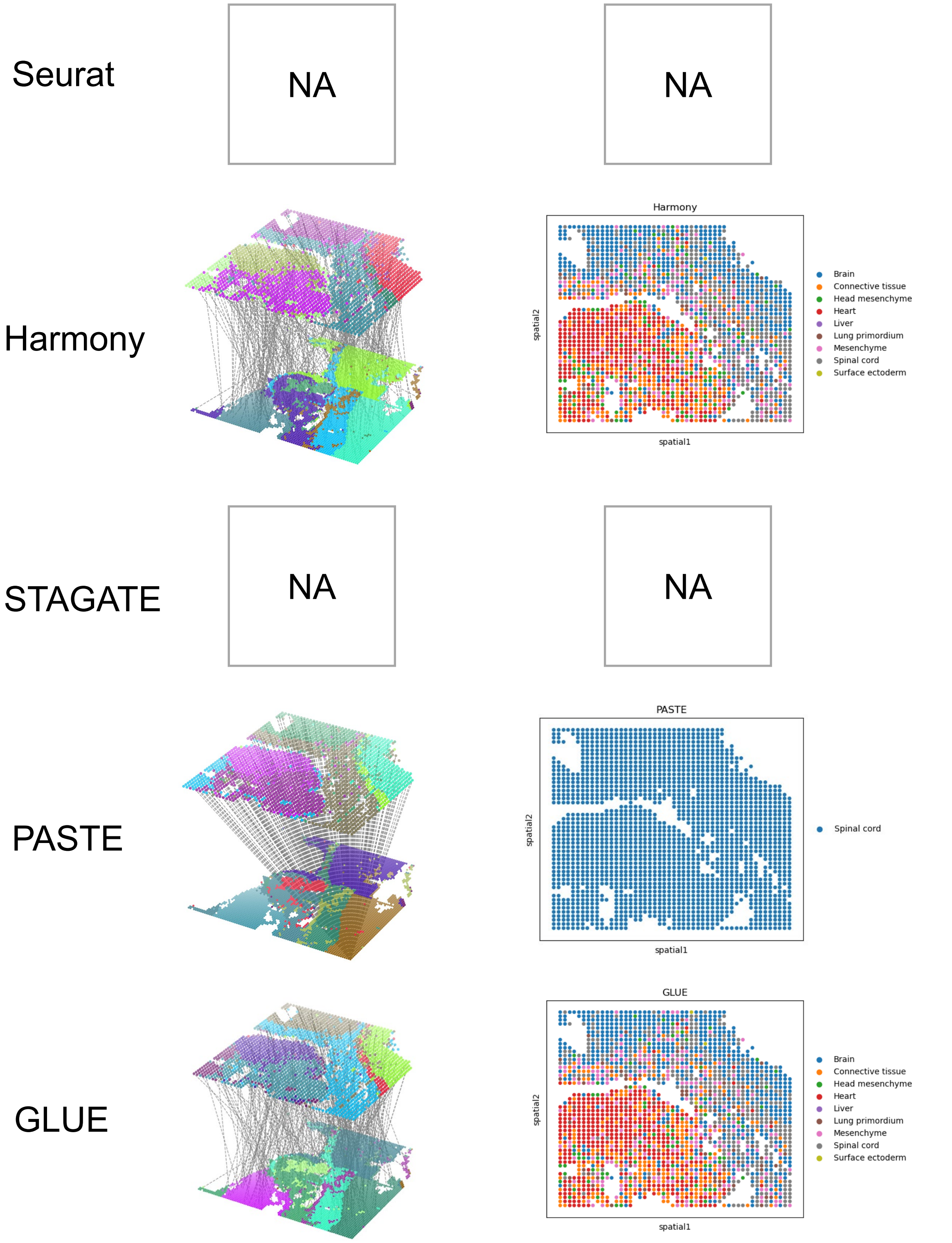


**Supplementary Fig. 7 Alignment results of current methods on the spatial-ATAC-seq and Stereo-seq slices.**

Visualization of the alignment results of spatial-ATAC-seq and Stereo-seq E11.5 mouse embryo slices via current methods based on GLUE embedding. The left panels show the complete alignment results (subsampled to 300 alignment pairs for clear visualization), while the right panels show cell type transfer results from Stereo-seq to spatial-ATAC-seq. Seurat and STAGATE failed because they do not support GLUE embedding as input.

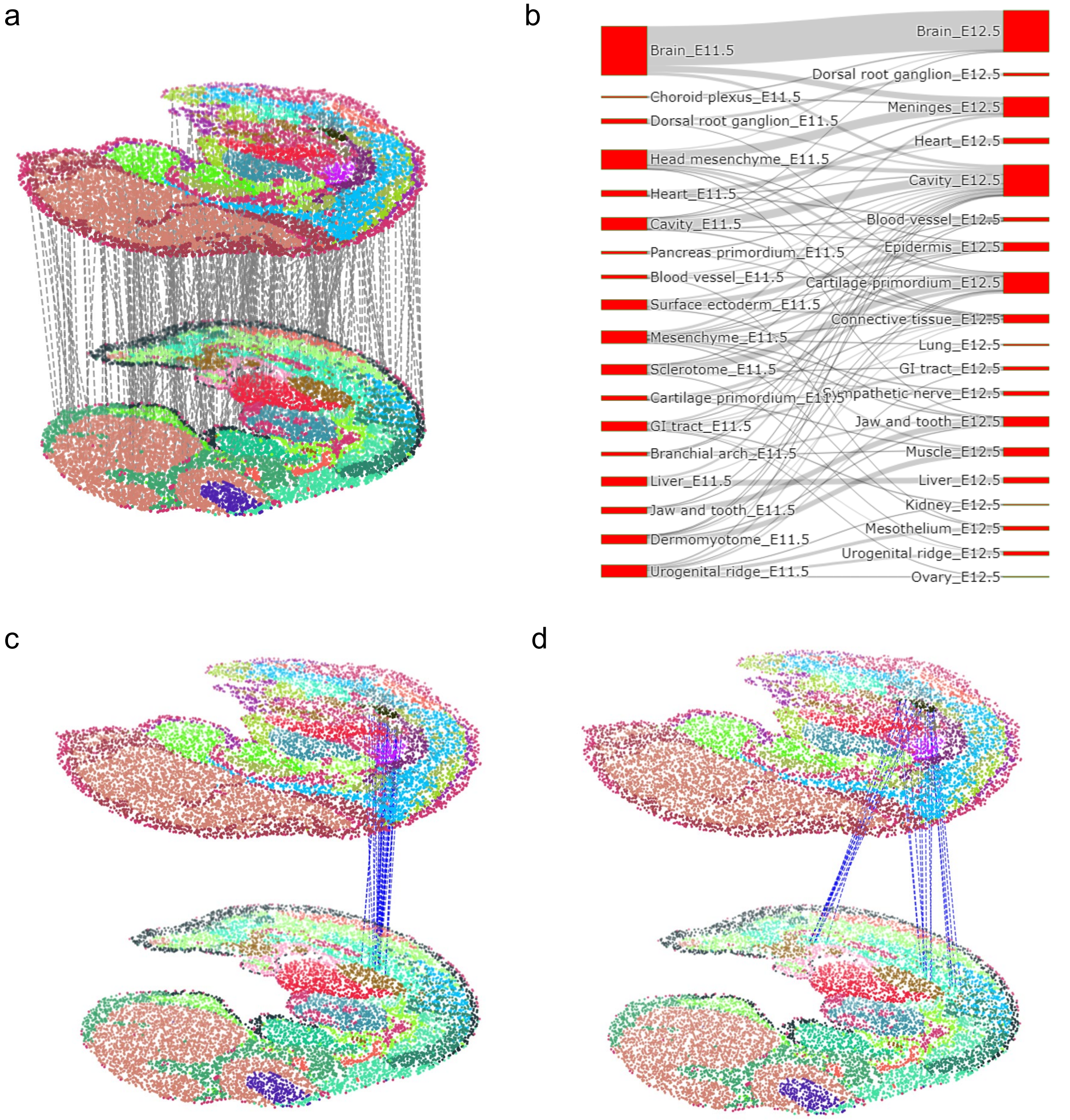


**Supplementary Fig. 8 Alignment of another mouse embryo E11.5 and E12.5 slices**
**a**,Visualization of the alignment results of another mouse embryo E11.5 and E12.5 mouse embryo slices. **b**, Sanky plot showing cell type correspondence of SLAT alignment between the two slices. **c**, **d**, Alignment visualization highlighting cells labeled as “Kidney” (**c**) and “Ovary” (**d**) in E12.5 and their aligned cells in E11.5, respectively.

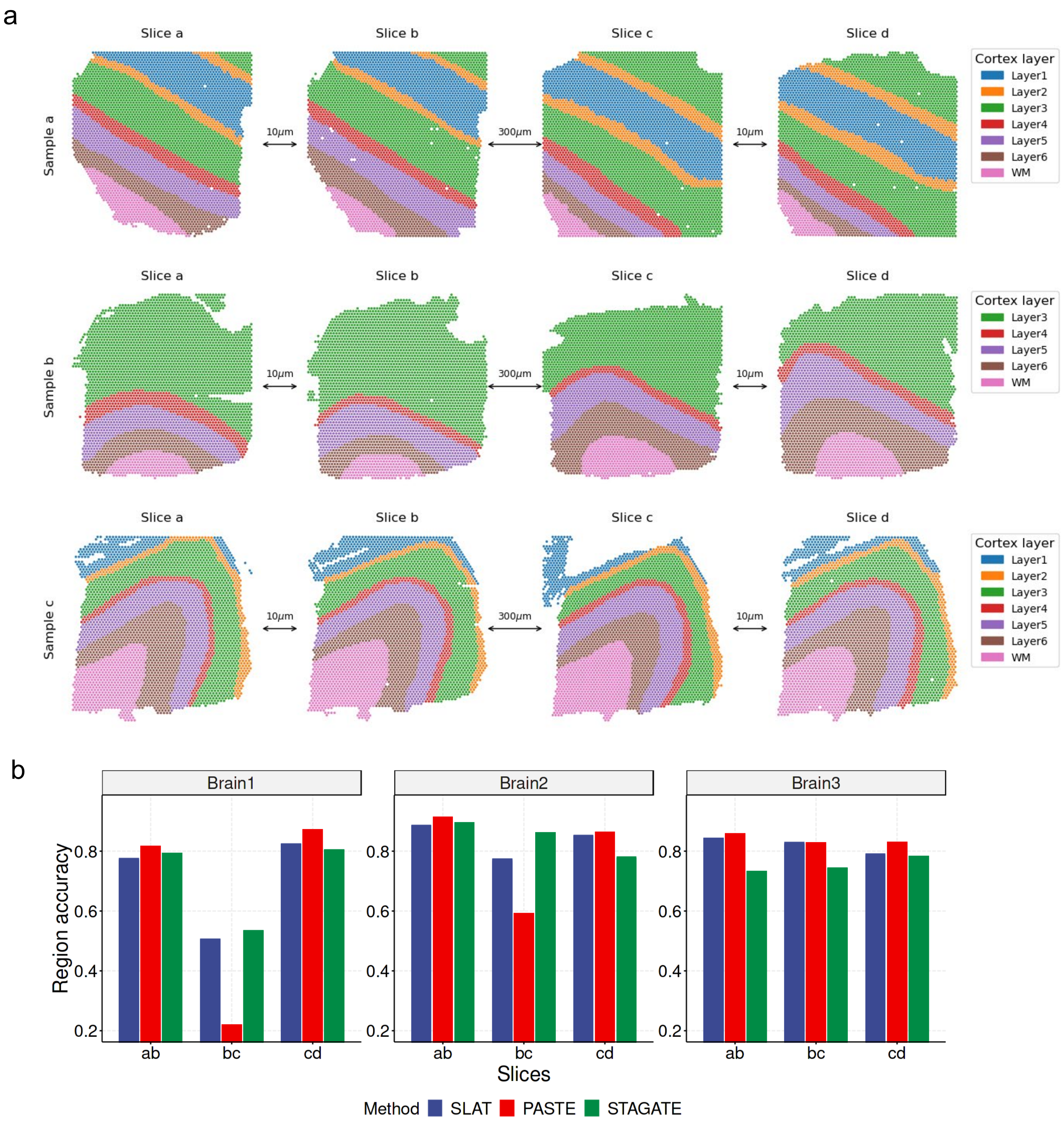


**Supplementary Fig. 9 Evaluation of 3D reconstruction on consecutive brain slices.** **a**, Visualization of 10x Visium slices used in 3D reconstruction benchmark (same 10x Visium datasets used in PASTE). Slices are divided into three groups, each group was consecutively sliced from the same human dorsolateral prefrontal cortex sample. The slices are colored by spatial regions annotated by original authors. **b**, Region matching accuracy[14] of different methods in 3D reconstruction. n = 8 repeats with different model random seeds. Dataset contains three groups, each with three consecutive slices (a, b and c, more details are in **Methods**).
