## Supplementary-Table for "Spatial-linked alignment tool (SLAT) for aligning heterogenous slices properly"

| Index | Publication | Species | Tissue | Technology | Resolution | Cells/Spots | Genes | Download Link |
|---|---|---|---|---|---|---|---|---|
| 1 | Chen et al. | Mouse | Whole embryo | Stereo-seq | 0.2μm | 5000-100,000 | >20,000 | https://db.cngb.org/stomics/mosta/download/ |
| 2 | Lohoff et al. | Mouse | Whole embryo | seqFISH | subcellular | ~10,000 | 351 | https://marionilab.cruk.cam.ac.uk/SpatialMouseAtlas/ |
| 3 | Deng et al. | Mouse | Whole embryo | spatial-ATAC-seq | 20μm | 2099 | >20,000 | https://www.ncbi.nlm.nih.gov/geo/query/acc.cgi?acc=GSE171943 |
| 4 | Jeffrey et al. | Mouse | Brain(hypothalamic preoptic) | MERFISH | subcellular | ~6,500 | 151 | https://datadryad.org/stash/dataset/doi:10.5061/dryad.8t8s248 |
| 5 | Kristen et al. | Human | Brain(dorsolateral prefrontal cortex, DLPFC) | 10x Visium | 50μm | ~3500 | >20,000 | https://github.com/LieberInstitute/spatialLIBD |
